# 4EBP1/2 support tumorigenicity and cell survival during energetic stress by translationally regulating fatty acid synthesis

**DOI:** 10.1101/2022.09.09.507243

**Authors:** Tal Levy, Kai Voeltzke, Laura Hauffe, Khawla Alasad, Marteinn Snaebjörnsson, Ran Marciano, Katerina Scharov, Mélanie Planque, Kim Vriens, Stefan Christen, Cornelius M Funk, Christina Hassiepen, Alisa Kahler, Beate Heider, Daniel Picard, Jonathan KM Lim, Zuelal Bas, Katja Bendrin, Andres Vargas-Toscano, Ulf Kahlert, Marc Remke, Moshe Elkabets, Thomas GP Grünewald, Andreas S. Reichert, Sarah-Maria Fendt, Almut Schulze, Guido Reifenberger, Barak Rotblat, Gabriel Leprivier

**Affiliations:** Department of Life Sciences, Faculty of Natural Sciences, Ben-Gurion University of the Negev, Beer-Sheva 84105, Israel; The National Institute for Biotechnology in the Negev, Ben-Gurion University of the Negev, Beer-Sheva 84105, Israel; Institute of Neuropathology, University Hospital Düsseldorf and Medical Faculty, Heinrich Heine University, 40225 Düsseldorf, Germany; Biochemistry and Molecular Biology, Theodor-Boveri-Institute, 97074 Würzburg, Germany; Division of Tumor Metabolism and Microenvironment, German Cancer Research Center (DKFZ), 69120 Heidelberg, Germany; Department of Pediatric Oncology, Hematology, and Clinical Immunology, University Hospital Düsseldorf and Medical Faculty, Heinrich Heine University, 40225 Düsseldorf, Germany; Laboratory of Cellular Metabolism and Metabolic Regulation, VIB-KU Leuven Center for Cancer Biology, VIB, 3000 Leuven, Belgium; Laboratory of Cellular Metabolism and Metabolic Regulation, Department of Oncology, KU Leuven and Leuven Cancer Institute (LKI), 3000 Leuven, Belgium; Division of Translational Pediatric Sarcoma Research, DKFZ, 69120 Heidelberg, Germany; Hopp Children’s Cancer Center (KiTZ), 69120 Heidelberg, Germany; German cancer consortium (DKTK) partner site Essen/Düsseldorf, 40225 Düsseldorf, Germany; Institute of Biochemistry and Molecular Biology I, Medical Faculty, Heinrich Heine University, 40225 Düsseldorf, Germany; Clinic for Neurosurgery, University Hospital Düsseldorf and Medical Faculty, Heinrich Heine University, 40225 Düsseldorf, Germany; Experimental and Clinical Research Center, Max-Delbrück Center for Molecular Medicine and Charité – Universitätsmedizin Berlin, corporate member of Freie Universität Berlin and Humboldt-Universität zu Berlin, 13125 Berlin, Germany; NeuroCure Clinical Research Center, Charité – Universitätsmedizin Berlin, corporate member of Freie Universität Berlin and Humboldt-Universität zu Berlin, 10117 Berlin, Germany; Molecular and Experimental Surgery, University Clinic for General-, Visceral, Vascular- and Transplantation Surgery, Faculty of Medicine and University Medicine, Otto-von-Guericke-University, 39120 Magdeburg, Germany; The Shraga Segal Department of Microbiology, Immunology and Genetics, Faculty of Health Science, Ben-Gurion University of the Negev, Beer-Sheva 84105, Israel; Faculty of Health Sciences, Ben-Gurion University of the Negev, Beer-Sheva 84105, Israel; Institute of Pathology, University Hospital Heidelberg, 69120 Heidelberg, Germany

**Keywords:** energetic stress, 4EBP1, mRNA translation, tumorigenesis

## Abstract

Energetic stress compels cells to evolve adaptive mechanisms to maintain homeostasis. Here, we report that the negative regulators of mRNA translation initiation eukaryotic initiation factor 4E binding proteins 1/2 (4EBP1/2) are essential to promote the survival of mammalian cells and budding yeast under glucose starvation. Functionally, 4EBP1/2 inhibit fatty acid synthesis upon energetic stress via repression of *Acetyl-CoA Carboxylase Alpha* (*ACACA*) mRNA translation, sparing NADPH, to maintain intracellular redox balance. This has important relevance in cancers, as we uncovered that oncogene-transformed cells and glioma cells exploit the 4EBP1/2 regulation of *ACACA* expression and redox balance to combat energetic stress, thereby supporting transformation and tumorigenicity in vitro and in vivo. Clinically, high *EIF4EBP1* (encoding 4EBP1) expression is associated with poor outcomes in several cancer types, including glioma. Our data reveal that 4EBP1/2 are conserved mediators of the survival response to energetic stress which are exploited by cancer cells for metabolic adaptation.

## INTRODUCTION

Glucose is one of the most important nutrients for living organisms. Lack of glucose has a profound biological impact on the cell. In order to adapt to glucose deprivation, cells have evolved highly coordinated and conserved metabolic responses. When glucose levels drop, anabolic processes, such as protein and fatty acid synthesis, are inhibited while catabolic processes, such as autophagy and fatty acid oxidation, are activated through the coordinated action of energy sensors, such as AMP-activated protein kinase (AMPK) (Lin and Hardie, 2018; Trefts and Shaw, 2021). Together, these responses preserve cellular energy and redox balance, thus maintaining cellular homeostasis (Caro-Maldonado, 2011).

A well-characterized pathological condition of glucose deprivation occurs in cancer (Gullino et al., 1964). Cancer cells growing within solid tumors experience glucose deprivation due to high glucose consumption combined with defects in tumor vasculature (Nagy et al., 2009; Sullivan et al., 2019). Matrix detachment, a hallmark of transformation and tumorigenicity, triggers glucose starvation-like energetic stress, characterized by ATP depletion and elevated reactive oxygen species (Schafer et al., 2009). Mechanisms promoting cell survival under matrix detachment also support cancer cell adaptation to glucose deprivation (Jeon et al., 2012). It is therefore important to delineate the mechanisms underlying metabolic adaptation to glucose deprivation, especially since this may reveal important actionable targets for therapeutic intervention in cancer (Hay, 2016).

One major cellular mechanism to preserve energy upon glucose deprivation is through inhibition of protein synthesis, the most energy consuming process in a cell (Buttgereit and Brand, 1995). Indeed, this process is inhibited under energy stress and failure to do so leads to cell death in both normal and cancer cells, which is assumed to be due to ATP depletion (Leprivier et al., 2015; Ng et al., 2012). However, it was reported that under energetic stress cells rather die from unresolved oxidative stress (Jeon et al., 2012; Schafer et al., 2009). Thus, it needs to be defined how the control of protein synthesis prevent cell death under energetic stress.

An important cellular energy sensor complex regulating protein synthesis is the mechanistic target of rapamycin complex 1 (mTORC1) (Leprivier and Rotblat, 2020; Orozco et al., 2020; Yoon et al., 2020). When energy is abundant, mTORC1 is active and promotes mRNA translation initiation by phosphorylating and activating 70-kDa ribosomal protein S6 kinase (p70S6K), and inhibiting eukaryotic initiation factor 4E binding proteins 1-3 (4EBPs). During energetic stress mTORC1 is inhibited, in turn releasing 4EBPs from inhibition, which leads to repression of mRNA translation initiation (Liu and Sabatini, 2020; Valvezan and Manning, 2019). It was reported that mTORC1 inhibition is absolutely required to preserve viability of normal and cancer cells under glucose starvation (Choo et al., 2010; Inoki et al., 2003). However, it is poorly understood how mTORC1 inhibition promotes cell survival during these conditions.

4EBPs are major repressors of protein synthesis regulated by glucose levels (Pause et al., 1994; Poulin et al., 1998). Since they were shown to promote the survival of *Drosophila* larvae challenged with nutrient starvation (Teleman et al., 2005; Tettweiler et al., 2005; Zid et al., 2009), these proteins may represent underappreciated regulators of the cellular and metabolic response to glucose starvation. In response to various stresses, 4EBPs inhibit mRNA translation initiation by binding eukaryotic initiation factor 4E (eIF4E), to prevent the formation of the eIF4E-containing pre-initiation complex required for cap-dependent mRNA translation (Silvera et al., 2010; Truitt and Ruggero, 2016). Recent findings highlight that 4EBPs preferentially inhibit the translation of hundreds of transcripts in a stress dependent manner, to regulate cellular processes such as proliferation, mitochondrial activity, and tumor cell invasion (Dowling et al., 2010; Hsieh et al., 2012; Morita et al., 2013; Thoreen et al., 2012). To date, the cellular functions of 4EBPs in eukaryotic cells challenged by energetic stress are not known, warranting further investigation. Moreover, such investigation would have profound implications for the role of 4EBPs in cancer, which is currently under debate (Musa et al., 2016). Namely, studies both support a tumor suppressive function of 4EBPs, as 4EBP1/2 double knock out (DKO) accelerates tumor development in genetically engineered mouse models of head and neck squamous cell carcinoma and prostate cancer (Ding et al., 2018; Wang et al., 2019), and a pro-tumorigenic function, with 4EBP1 supporting breast cancer progression in vivo (Braunstein et al., 2007).

Here, we found that 4EBP1/2 promote cell survival during glucose starvation in yeast, mouse, and human cells. Mechanistically, 4EBP1/2 curb fatty acid synthesis in response to glucose starvation, by selectively repressing the translation of *Acetyl-CoA Carboxylase Alpha* (*ACACA*) mRNA. This allows the preservation of NADPH levels, which are normally consumed by fatty acid synthesis, thereby preventing oxidative stress and cell death during glucose starvation. Furthermore, we report that 4EBP1/2 regulation of *ACACA* expression is exploited by cancer and transformed cells to support tumorigenicity in vitro and promote tumor growth in vivo. Finally, we reveal that 4EBP1 expression has clinical relevance in multiple cancers, and is functional in glioma to promote tumor growth and aggressiveness, altogether highlighting a pro-tumorigenic function for the mRNA translation inhibitor 4EBP1.

## RESULTS

### 4EBP1/2 promote cell survival under glucose deprivation by controlling mRNA translation

We first investigated the function of 4EBPs in the cellular response to glucose starvation. To do so, we assessed the impact of 4EBP1/2 depletion on cell survival during glucose starvation, using cells deficient for 4EBP activity, namely 4EBP1/2 DKO mouse embryonic fibroblasts (MEFs) or stable 4EBP1/2 knock down (kd) (sh4EBP1/2) HEK293 cells (Dowling et al., 2010). Glucose starvation led to massive cell death of 4EBP1/2 DKO MEFs as well as sh4EBP1/2 HEK293 cells compared to their control counterparts (WT and shScr, respectively), which was not the case in glucose-containing basal conditions (Fig. 1A&B). This was confirmed in cell lines of other origins, including induced pluripotent stem cells (iPSC), breast cancer cells (MCF7), neuroblastoma cells (IMR-32 and Kelly), and medulloblastoma cells (HD-MB03 and Med8a) (Fig. S1A-F). In these cells, stable kd of 4EBP1 severely restricted survival under glucose deprivation (Fig. S1A-F). Conversely, overexpression of a constitutively active 4EBP1 mutant 4EBP1 (T37A/T46A) (4EBP1^AA^) in HeLa cells, which are normally sensitive to nutrients withdrawal (Leprivier et al., 2013), was sufficient to protect them from the induction of cell death under glucose starvation (Fig. 1C). In contrast, manipulating 4EBP1/2 expression levels had no impact on cell death rates triggered by amino acid starvation (Fig. 1A-C), pointing toward a specific role of 4EBP1/2 in response to glucose starvation.

**Figure 1.**
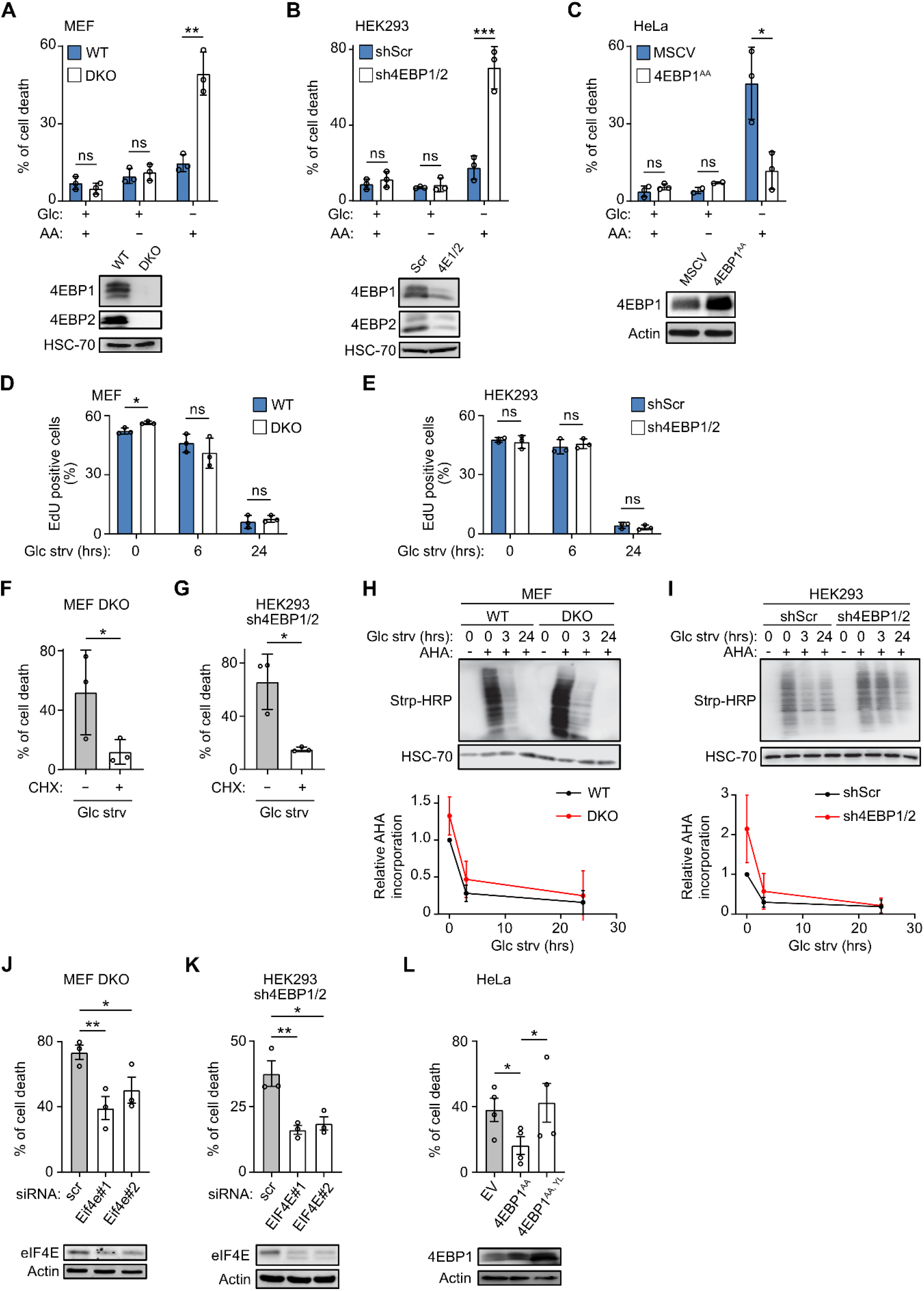
4EBP1/2 prevent cell death in response to glucose starvation through control of eIF4E and mRNA translation. (A-C) WT and 4EBP1/4EBP2 DKO MEF (A), control (shScr) and stable 4EBP1/4EBP2 knock down (sh4EBP1/2) HEK293 (B), or control (MSCV) and stable 4EBP1^AA^ overexpressing HeLa cells (C) were grown in complete media, or starved for amino acids (AA) or for glucose (Glc) for 48 hrs. Cell death was measured by propidium iodide (PI) staining and flow cytometry. The level of the indicated proteins was analyzed by immunoblotting. (D, E) WT and 4EBP1/4EBP2 DKO MEF (D), or shScr and sh4EBP1/2 HEK293 cells (E) were grown in complete medium or glucose starved (Glc strv) for the indicated times and labeled with EdU. The percentage of EdU positive cells was analyzed by flow cytometry. (F, G) 4EBP1/4EBP2 DKO MEF (F) or sh4EBP1/2 HEK293 cells (G) were grown in glucose starved medium (Glc strv) with or without cycloheximide (CHX) for 48 hrs. Cell death was analyzed as in (A-C). (H, I) WT and 4EBP1/4EBP2 DKO MEF (H), or shScr and sh4EBP1/2 HEK293 cells (I) were grown in complete medium or glucose starved (Glc strv) for the indicated times and labeled with azidohomoalanine (AHA). Levels of AHA-labelled proteins were detected by immunoblotting. (J, K) 4EBP1/4EBP2 DKO MEF (J) or sh4EBP1/2 HEK293 cells (K) were transfected with control siRNA (scr) or siRNAs targeting *EIF4E* and grown in glucose starved medium (Glc strv) for 48 hrs. Cell death and protein levels were analyzed as in (A-C). (L) HeLa cells stably expressing empty vector (EV), 4EBP1^AA^ or 4EBP1^AA,YL^ were grown in glucose starved medium (Glc strv) for 48 hrs. Cell death and protein levels were analyzed as in (A-C). Where shown, data are reported as means ± SD with indicated significance (*p < 0.05, **p < 0.01, and ***p < 0.005).

We next asked which cellular processes may mediate 4EBP1/2 protective function under glucose starvation. Since it was reported that 4EBP1/2 exerts control on cell proliferation upon pharmacological inhibition of mTORC1, amino acid starvation, or serum starvation (Dowling et al., 2010), we first tested the involvement of this cellular process. Interestingly, while proliferation rates were severely reduced following 24 hrs glucose deprivation in all cell lines tested, we observed no difference in proliferative capacity between wild type (WT) and 4EBP1/2 DKO MEFs, or between shScr and sh4EBP1/2 HEK293 cells (Fig. 1D&E). Similarly, we found no evidence that autophagy was responsible for the observed function of 4EBP1/2 under glucose starvation, as rates of autophagy were similar in control and 4EBP1/2 deficient MEFs and HEK293 cells in such conditions (Fig. S1G&H). We also verified that the loss of 4EBP1/2 did not prevent the activation of the energy sensor and pro-survival regulator AMPK in response to glucose deprivation (Fig. S1I&J).

In addition, since 4EBP1/2 are major repressors of mRNA translation initiation, we examined the contribution of mRNA translation activity to 4EBP1/2 protective function under glucose starvation. Notably, pharmacological inhibition of protein synthesis using cycloheximide (CHX) fully rescued 4EBP1/2 deficient MEFs and HEK293 cells from glucose starvation-induced cell death (Fig. 1F&G) and significantly reduced cell death in sh4EBP1 MCF7 cells (Fig. S1B).

This suggests that uncontrolled protein synthesis in cells lacking 4EBP1/2 may contribute to glucose starvation-induced cell death.

Unexpectedly, using azidohomoalanine (AHA) labeling and Click Chemistry (Marciano et al., 2018), we detected similarly low rates of overall protein synthesis under glucose starvation in control and 4EBP1/2 deficient MEFs and HEK293 cells (Fig. 1H&I). In contrast, upon pharmacological inhibition of mTORC1 (using KU-0063794), 4EBP1/2 deficient cells exhibited high rates of global protein synthesis vs. control cells (Fig. S1K), suggesting that 4EBPs protective function under glucose starvation is independent from the inhibition of global protein synthesis. Together, our data suggest that a selective, rather than global, regulation of mRNA translation by 4EBP1/2 promotes cell viability under glucose deprivation.

Given that 4EBP1/2 function by binding to eIF4E to selectively repress the translation of a subset of transcripts (Dowling et al., 2010; Hsieh et al., 2012; Morita et al., 2013; Thoreen et al., 2012), we tested the involvement of eIF4E in the protective function of 4EBP1/2. Notably, kd of eIF4E in both 4EBP1/2 DKO MEFs and sh4EBP1/2 HEK293 cells led to a significant reduction of cell death under glucose starvation (Fig. 1J&K). In addition, forced expression of an eIF4E-non-binding mutant of 4EBP1^AA^, 4EBP1 (Y54A/L59A) (4EBP1^AA, YL^; (Mader et al., 1995)), failed to prevent cell death of HeLa cells upon glucose depletion, in contrast to 4EBP1^AA^ (Fig. 1L), indicating that binding of 4EBP1/2 to eIF4E is required for 4EBP1/2-mediated cellular protection under glucose-deprived conditions. Therefore, 4EBP1/2 exert a pro-survival function under glucose deprivation by binding and inhibiting eIF4E to regulate mRNA translation.

### 4EBP1/2 maintain redox balance to preserve cell viability under glucose deprivation

To dissect how 4EBP1/2 protect cells under glucose-deprived conditions, we assessed the impact of 4EBP1/2 on cellular energetic and redox balances, which are major cellular parameters influenced by glucose availability (Hay, 2016; Jeon et al., 2012). While ATP levels were reduced following glucose starvation as anticipated, 4EBP1/2 deficient MEFs and HEK293 cells showed the same amounts of ATP as compared to corresponding control cells under these conditions (Fig. S2A&B). Given that protein synthesis is the most ATP-consuming process within a cell (Buttgereit and Brand, 1995), this result is consistent with the lack of impact of 4EBP1/2 on rates of overall protein synthesis under these conditions (Fig. 1H&I).

In terms of redox balance, endogenous H_2_O_2_ levels were higher under glucose deprivation in 4EBP1/2 DKO MEFs and sh4EBP1/2 HEK293 cells compared to their respective controls (WT and shScr, respectively) (Fig. 2A&B). In addition, overexpression of 4EBP1^AA^ prevented such increases of H_2_O_2_ levels in HeLa cells during glucose depletion (Fig. 2C). These findings suggest a role for 4EBP1/2 in controlling redox balance upon glucose starvation.

**Figure 2.**
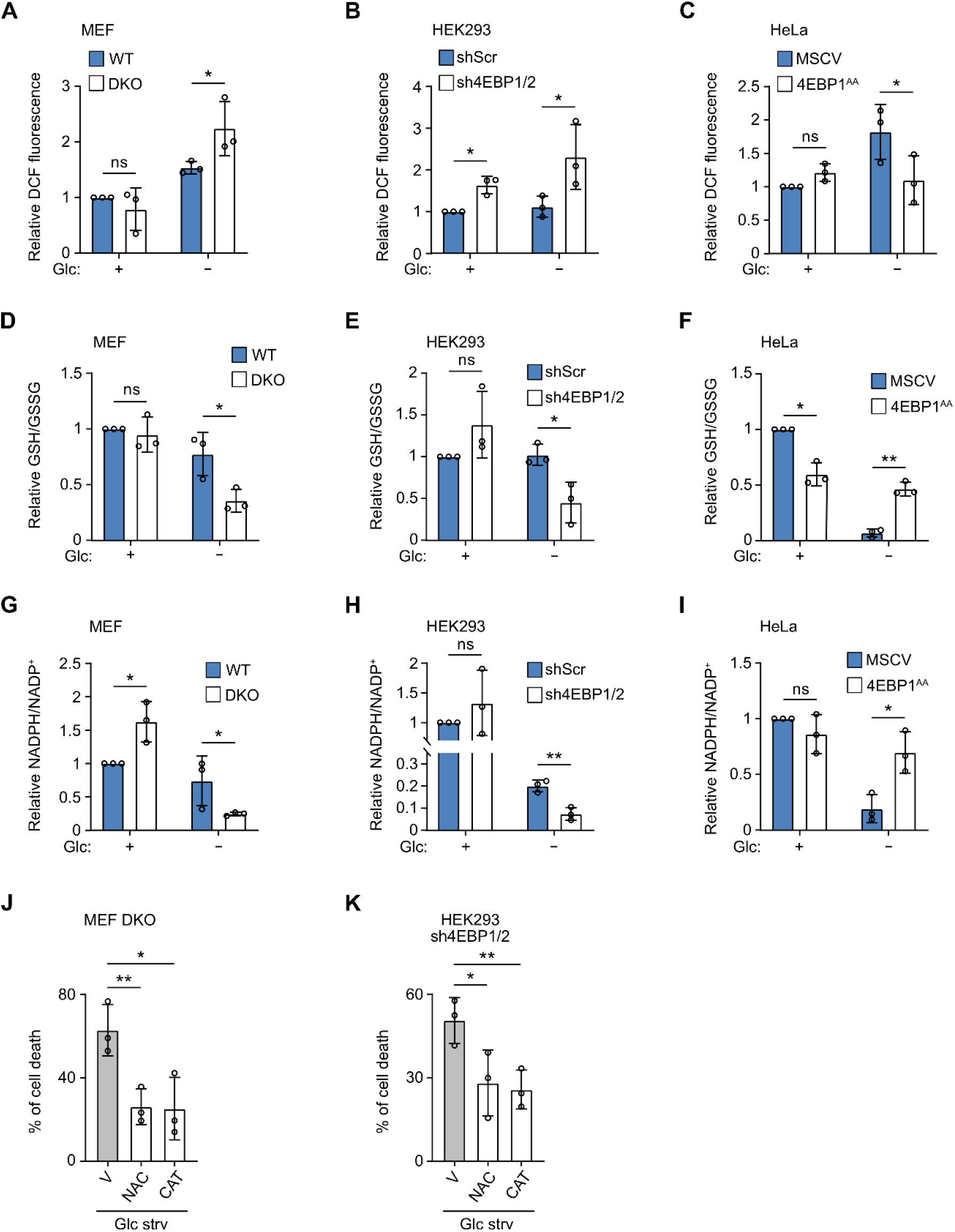
4EBP1/2 preserve the redox balance under glucose starvation by maintaining antioxidant power. (A-C) WT and 4EBP1/4EBP2 DKO MEF (A), shScr and sh4EBP1/2 HEK293 (B), or MSCV and stable 4EBP1^AA^ overexpressing HeLa cells (C) grown in complete medium or glucose (Glc) starved for 24 hrs were labelled with CM-DCFDA and analyzed by flow cytometry. (D-F) WT and 4EBP1/4EBP2 DKO MEF (D), shScr and sh4EBP1/2 HEK293 (E), or MSCV and stable 4EBP1^AA^ overexpressing HeLa cells (F) were grown in complete medium or glucose (Glc) starved for 24 hrs, and reduced and total glutathione were measured and expressed as the ratio of reduced (GSH) to oxidized (GSSG) glutathione. (G-I) WT and 4EBP1/4EBP2 DKO MEF (G), shScr and sh4EBP1/2 HEK293 (H), or MSCV and stable 4EBP1^AA^ overexpressing HeLa cells (I) were grown in complete medium or glucose (Glc) starved for 24 hrs, and NADP^+^ and NADPH levels were measured. (J, K) 4EBP1/4EBP2 DKO MEF (J) or sh4EBP1/2 HEK293 cells (K) were grown in glucose starved medium (Glc strv) and treated with vehicle (V), N-acetyl cysteine (NAC) or Catalase (CAT) for 48 hrs. Cell death was measured by PI staining and flow cytometry. Where shown, data are reported as means ± SD with indicated significance (*p < 0.05, and **p < 0.01).

Glutathione is a major cellular antioxidant, and cells recycle the oxidized form, GSSG, to the reduced form, GSH, making the GSH to GSSG ratio an indicator of oxidative stress (Flohe, 2013). In line with this, 4EBP1/2 DKO MEFs and sh4EBP1/2 HEK293 cells exhibited a lower ratio of reduced to oxidized glutathione (GSH/GSSG), relative to their respective control cells under glucose starvation, indicative of lower antioxidant capacity in 4EBP1/2 deficient cells (Fig. 2D&E). Conversely, 4EBP1^AA^ overexpression precluded severe depletion of GSH/GSSG ratio in HeLa cells upon glucose removal (Fig. 2F). Since we did not observe changes in the level of total glutathione in 4EBP1/2 deficient MEFs and HEK293 or in 4EBP1^AA^ overexpressing cells during glucose deprivation (Fig. S2C-E), we reasoned that in glucose-starved cells, 4EBP1/2 may contribute to the recycling of oxidized glutathione (GSSG) to its reduced form (GSH), rather than its biosynthesis.

The conversion of GSSG to GSH requires the oxidation of NADPH to NADP^+^, whose levels are a strong determinant of cellular survival during glucose deprivation (Jeon et al., 2012). Notably, 4EBP1/2 DKO MEFs and sh4EBP1/2 HEK293 cells were characterized by acute decrease of NADPH/NADP^+^ ratio under glucose withdrawal compared with respective control cells (Fig. 2G&H). Furthermore, 4EBP1^AA^ overexpression led to a significant increase in the NADPH/NADP^+^ ratio in HeLa cells during glucose starvation (Fig. 2I). These results are consistent with a role of 4EBP1/2 in promoting NADPH levels under glucose depletion.

We next asked whether increased oxidative stress, observed in glucose-starved, 4EBP1/2 deficient cells, is linked to increased sensitivity to glucose starvation. To address this question, we supplemented glucose-starved 4EBP1/2 DKO MEFs and sh4EBP1/2 HEK293 cells with antioxidants, namely N-acetyl cysteine (NAC) or Catalase (CAT), and found that these antioxidants significantly reduced cell death compared to vehicle-treated cells (Fig. 2J&K). We conclude that the protective function of 4EBP1/2 under glucose deprivation relies on curbing oxidative stress by promoting NADPH levels.

### 4EBP1/2 control fatty acid synthesis under glucose deprivation to preserve NADPH levels

We next asked how 4EBP1/2 support NADPH levels under glucose deprivation. We focused our attention on fatty acid synthesis, the most NADPH-consuming process within a cell (Fan et al., 2014). Importantly, to survive glucose starvation, cells must inhibit fatty acid synthesis to preserve NADPH levels (Jeon et al., 2012). Therefore, we asked whether 4EBP1/2 contribute to the inhibition of fatty acid synthesis induced by glucose starvation. Using ^14^C acetate labeling, we measured the impact of 4EBP1/2 on fatty acid synthesis by quantifying ^14^C incorporation into the cellular lipid fraction under basal conditions and after 24 hrs of glucose starvation (Fig. 3A). As expected, we found that glucose starvation led to reduced ^14^C-labelled lipids in 4EBP1/2 WT MEFs and shScr HEK293 cells (Fig. 3B&C). This was not observed in 4EBP1/2 DKO MEFs and sh4EBP1/2 HEK293 cells which displayed higher amounts of ^14^C-labelled lipids than corresponding control cells under glucose starvation (Fig. 3B&C). Altogether, these data suggest that 4EBP1/2 are essential for cells to inhibit fatty acid synthesis activity in response to glucose starvation.

**Figure 3.**
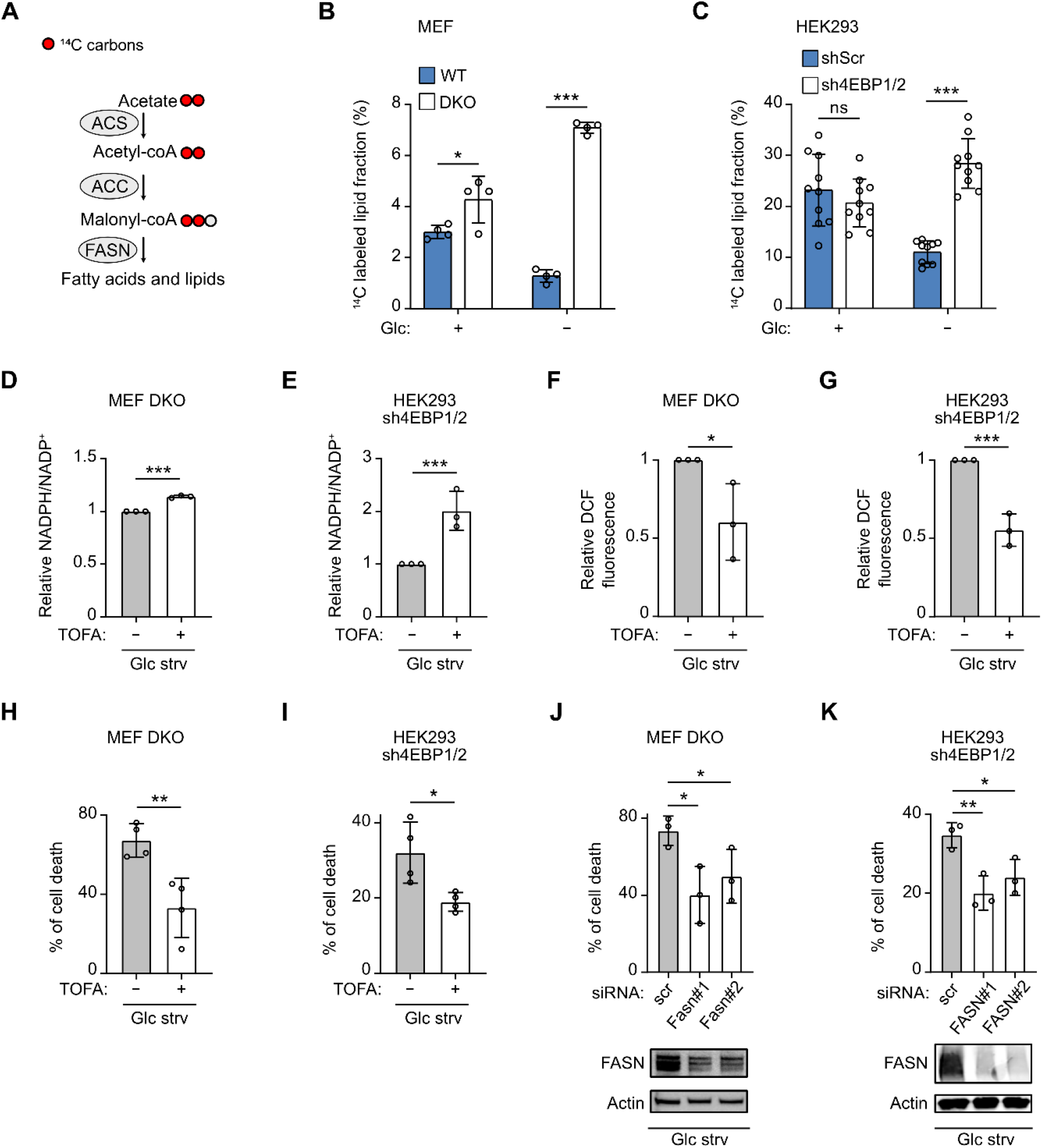
4EBP1/2 control fatty acid synthesis activity in response to glucose starvation to preserve redox balance and protects cells. (A) Scheme of the [^14^C] acetate labeling assay to measure fatty acid synthesis activity. ACS: acetyl-CoA synthetase, ACC: acetyl-CoA carboxylase, FASN: fatty acid synthase. (B, C) WT and 4EBP1/4EBP2 DKO MEF (B), or shScr and sh4EBP1/2 HEK293 cells (C) grown in complete medium or glucose (Glc) starved for 24 hrs were labeled with [^14^C] acetate in the last 18 hrs. [^14^C] was measured in the lipid fraction using a scintillation counter and normalized to total protein levels. (D, E) 4EBP1/4EBP2 DKO MEF (D) or sh4EBP1/2 HEK293 cells (E) were grown in glucose starved medium (Glc strv) for 24 hrs with or without TOFA, and NADP^+^ and NADPH levels were measured. (F, G) 4EBP1/4EBP2 DKO MEF (F) or sh4EBP1/2 HEK293 cells (G) grown in glucose starved medium (Glc strv) for 24 hrs with or without TOFA were labelled with CM-DCFDA and analyzed by flow cytometry. (H, I) 4EBP1/4EBP2 DKO MEF (H) or sh4EBP1/2 HEK293 cells (I) were grown in glucose starved medium (Glc strv) with or without TOFA for 48 hrs. Cell death was measured by PI staining and flow cytometry. (J, K) 4EBP1/4EBP2 DKO MEF (J) or sh4EBP1/2 HEK293 cells (K) were transfected with control siRNA (scr) or siRNAs targeting *FASN* and were grown in glucose starved medium (Glc strv) for 48 hrs. Cell death was analyzed as in (H, I). FASN protein levels were analyzed by immunoblotting. Where shown, data are reported as means ± SD with indicated significance (*p < 0.05, **p < 0.01, and ***p < 0.005).

We next wondered whether increased fatty acid synthesis is responsible for the decreased NADPH levels and increased ROS observed in glucose-starved 4EBP1/2 deficient cells. To address this question, we pharmacologically inhibited fatty acid synthesis in 4EBP1/2 DKO MEFs and sh4EBP1/2 HEK293 cells using the acetyl-CoA carboxylase (ACC) inhibitor TOFA. This led to a significant elevation of the NADPH/NADP^+^ ratio (Fig. 3D&E) and reduced ROS levels in these cells upon glucose withdrawal (Fig. 3F&G), thus supporting that 4EBP1/2 control redox balance under glucose-limited conditions by curbing fatty acid synthesis.

Given that fatty acid synthesis activity is a determinant of cell viability under glucose starvation, we next assessed whether increased fatty acid synthesis contributes to the sensitivity of 4EBP1/2-depleted cells under glucose starvation. To this end, we treated 4EBP1/2 deficient MEFs and HEK293 cells with TOFA, which was able to rescue glucose-deprived cells from death (Fig. 3H&I). This was also confirmed by selectively targeting fatty acid synthase (FASN), the enzyme consuming NADPH during fatty acid synthesis. Namely, siRNA-mediated kd of *FASN* in 4EBP1/2 DKO MEFs and sh4EBP1/2 HEK293 cells significantly reduced the levels of glucose withdrawal-induced cell death, as compared to control siRNA (scr) (Fig. 3J&K). Altogether, our findings support that during glucose starvation, 4EBP1/2 promote cell viability by inhibiting fatty acid synthesis to preserve NADPH levels and reduce ROS.

### 4EBP1/2 selectively regulate the translation of *ACACA* to preserve cell viability under glucose deprivation

We next sought to determine the mechanism by which 4EBP1/2 regulate fatty acid synthesis in response to glucose starvation. Since 4EBP1/2 were reported to selectively block the translation of specific transcripts, we asked whether 4EBP1/2 could restrict the synthesis of one of the fatty acid synthesis enzymes. Immunoblot analysis showed that levels of ATP citrate lyase (ACLY), ACC2 and FASN were similar in control and 4EBP1/2 DKO MEFs or sh4EBP1/2 HEK293 cells under glucose starvation (Fig. 4A&B). In contrast, expression of fatty acid synthesis rate-limiting enzyme ACC1 was severely decreased in control MEFs and HEK293 cells by glucose depletion, which was not the case in the corresponding 4EBP1/2 deficient cells, thus revealing that ACC1 is still highly expressed in 4EBP1/2 deficient cells under glucose starvation (Fig. 4A&B). In agreement, 4EBP1^AA^ overexpression led to a reduction of ACC1 levels in HeLa cells under glucose deprivation (Fig. S3A). Intriguingly, levels of inactive, phosphorylated ACC1 were higher in 4EBP1/2 deficient versus control cells under glucose deprivation (Fig. 4A&B), similarly to ACC1 expression pattern. The difference in ACC1 expression between control and 4EBP1/2 deficient HEK293 cells under glucose deprivation was not due to changes in *ACACA* (gene encoding ACC1) mRNA level (Fig. 4C), suggesting regulation at the translational level. To ascertain whether 4EBP1/2 preferentially control *ACACA* translation under glucose starvation, we quantified levels of *ACACA* transcripts in polysomal and total mRNA and calculated the translation efficiency as the ratio of polysomal to total mRNA levels (Liang et al., 2018). We found that translation efficiency of *ACACA* transcript was significantly higher in sh4EBP1/2 HEK293 cells compared to control cells under glucose deprivation (Fig. 4D).

**Figure 4.**
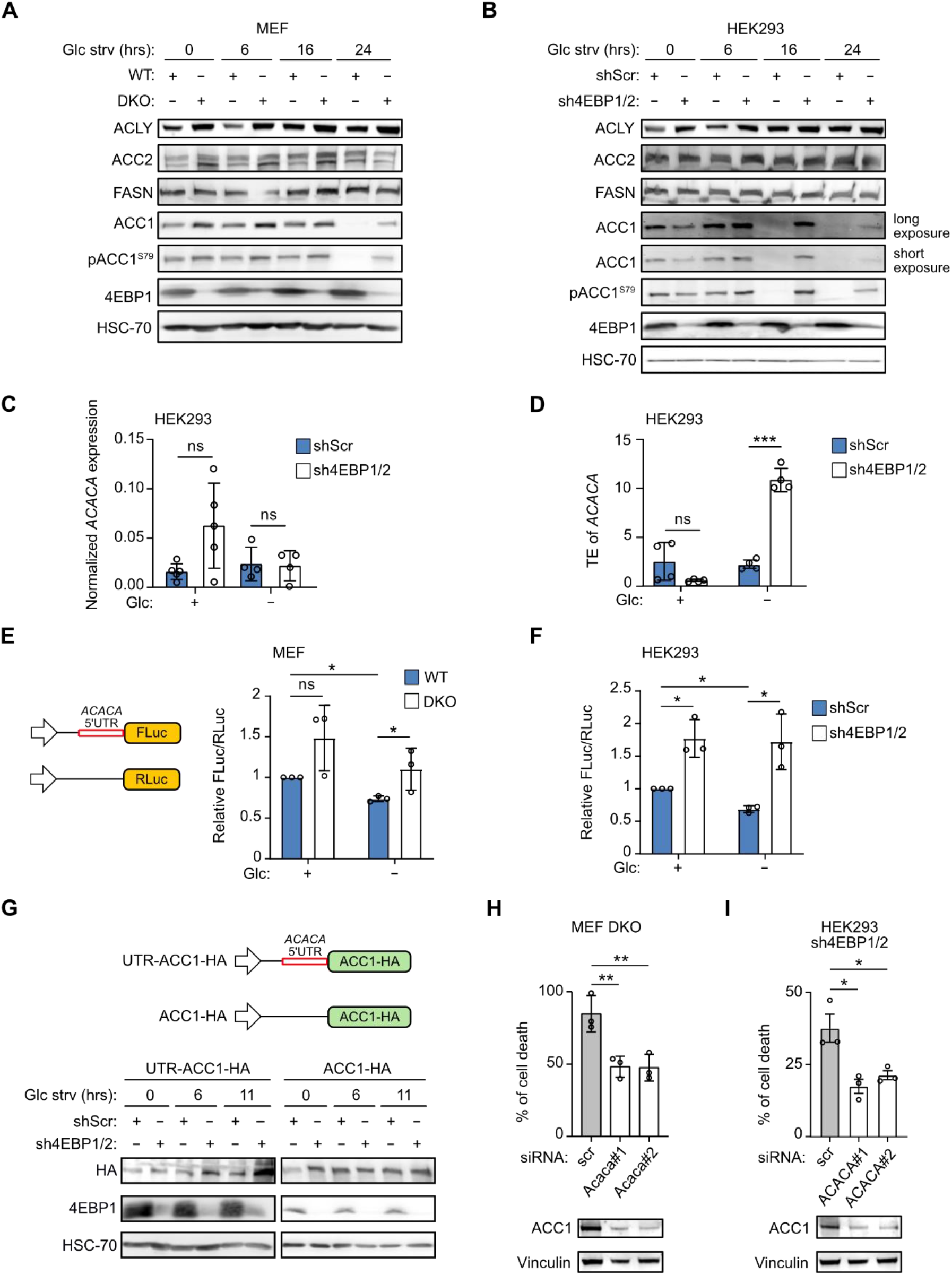
4EBP1/2 repress *ACACA* translation under glucose starvation. (A, B) WT and 4EBP1/4EBP2 DKO MEF (A), or shScr and sh4EBP1/2 HEK293 cells (B) were grown in complete medium or glucose starved (Glc strv) for the indicated times, and analyzed by immunoblotting using antibodies against the indicated proteins. (C) ShScr and sh4EBP1/2 HEK293 cells were grown in complete medium or glucose starved (Glc strv) for 16 hrs, and *ACACA* mRNA expression was analyzed by qRT-PCR. (D) ShScr and sh4EBP1/2 HEK293 cells were grown in complete medium or glucose starved (Glc strv) for 6 hrs, and translation efficiency (TE) of *ACACA* mRNA was calculated by measuring the levels of polysomal and total *ACACA* mRNA by qRT-PCR. (E, F) WT and 4EBP1/4EBP2 DKO MEF (E), or shScr and sh4EBP1/2 HEK293 cells (F) were transfected with an *ACACA* 5’UTR-containing Firefly Luciferase construct and a control *Renilla* Luciferase vector. Cells were grown in complete medium or glucose starved (Glc strv) for 6 hrs, and luminescence was measured. (G) 293 cells were transfected with an HA-tagged ACC1 expressing vector with or without the *ACACA* 5’UTR. Cells were grown in complete medium or glucose starved (Glc strv) for the indicated times, and analyzed by immunoblotting using antibodies against the indicated proteins. (H, I) 4EBP1/4EBP2 DKO MEF (H) or sh4EBP1/2 HEK293 cells (I) were transfected with control siRNA (scr) or siRNAs targeting *ACACA* and grown in glucose starved medium (Glc strv) for 48 hrs. Cell death was measured by PI staining and flow cytometry. ACC1 protein levels were analyzed by immunoblotting. Where shown, data are reported as means ± SD with indicated significance (*p < 0.05, **p < 0.01, and ***p < 0.005).

Given that 4EBP/eIF4E-mediated selective translational control is mediated through each target’s 5’UTR, we investigated the potential regulation of *ACACA* 5’UTR by 4EBP1/2 in response to glucose starvation. Since *ACACA* encodes several isoforms harboring different 5’UTR (Damiano et al., 2018), we focused on the 5’UTR present in human *ACACA* transcript variant 3, which is also highly conserved in mice. Notably, we observed that *ACACA* 5’UTR activity, as monitored with a luciferase reporter, was significantly decreased upon glucose starvation in control MEFs and HEK293 cells (Fig. 4E&F). More importantly, *ACACA* 5’UTR activity was higher in 4EBP1/2 DKO MEFs and sh4EBP1/2 HEK293 cells under glucose starvation compared to respective control cells (Fig. 4E&F).

To determine the contribution of the 5’UTR to 4EBP1/2 regulation of ACC1 expression in the context of the *ACACA* transcript, we ectopically expressed HA-tagged *ACACA*, flanked or not by the 5’UTR (Fig. 4G), and monitored the impact of 4EBP1/2 on exogenous ACC1 protein levels during glucose starvation. While the expression of ACC1 with no 5’UTR did not differ between sh4EBP1/2 and control HEK293 cells under glucose starvation, the level of 5’UTR-containing ACC1 was higher in sh4EBP1/2 cells as compared to controls during glucose starvation (Fig. 4G). Altogether, these data support that *ACACA* 5’UTR plays a role in 4EBP1/2-mediated inhibition of *ACACA* translation under glucose-deprived conditions. To determine the contribution of increased ACC1 expression towards the sensitivity of 4EBP1/2 DKOs and sh4EBP1/2 HEK293 cells to glucose starvation, we assessed the impact of ACC1 kd on cell viability during glucose starvation. We found that ACC1 kd rescued 4EBP1/2 deficient cells from glucose starvation-induced cell death (Fig. 4H&I), in line with our findings using an ACC inhibitor (Fig. 3H&I). These results highlight that 4EBP1/2 protect cells from glucose starvation by inhibiting the translation of *ACACA* in a 5’UTR dependent manner, thus reducing fatty acid synthesis, preserving NADPH, and limiting oxidative stress.

### The 4EBP1/2 orthologue Eap1 preserves viability of yeast under glucose deprivation

Since most living organisms need to cope with glucose-starved conditions, we asked whether the 4EBP1/2-mediated protective function under glucose starvation represents an evolutionarily conserved biological response to such a stress. We aimed to delineate the function of 4EBP under glucose deprivation in an evolutionary distant model organism, *Saccharomyces cerevisiae*, by studying the two functional 4EBP orthologues in yeast Eap1p and Caf20p (Cosentino et al., 2000; Lanker et al., 1992). Disruption of either *eap1* (*eap1Δ*) or *caf20* (*caf20Δ*) had no observable impact on the growth rate of serially diluted yeast cultures on agar in comparison to the WT strain grown in glucose-containing rich YPD medium (Fig. 5A), which is in agreement with previous reports stating that *EAP1* and *CAF20* are not essential genes (Cosentino et al., 2000; Lanker et al., 1992). However, the growth of *eap1Δ* strain, but not of *caf20Δ* strain, was severely compromised on solid rich glucose-free YP media, as compared to WT strain (Fig. 5A) and the deletion of both (*eap1Δ*/*caf20Δ*) had no additional detrimental impact on growth in glucose-free medium compared to *eap1Δ* Strain alone (Fig. 5A). Moreover, we confirmed that in liquid media, *eap1* disruption had a pronounced negative effect on growth in glucose-free YP medium compared to WT strain (Fig. 5B).

We next assessed whether such a phenotype was due to a reduction in survival under glucose-deprived conditions. Clonogenic assays of WT and *eap1Δ* strains grown in liquid conditions in the absence of glucose indicated that disruption of *eap1* prevented survival of yeast upon glucose withdrawal (Fig. 5C). Overall, our data indicate that the 4EBP orthologue Eap1p, but not Caf20p, promotes survival of *Saccharomyces cerevisiae* in glucose-deprived conditions, thus supporting the premise that 4EBP1/2 protective function in glucose starvation is evolutionarily conserved. The functional difference between Eap1p and Caf20p is in line with differential requirements of these proteins for growth under varying forms of nutrient-depleted conditions and may arise from differences in the set of transcripts translationally regulated by Eap1p and Caf20p (Cridge et al., 2010).

**Figure 5.**
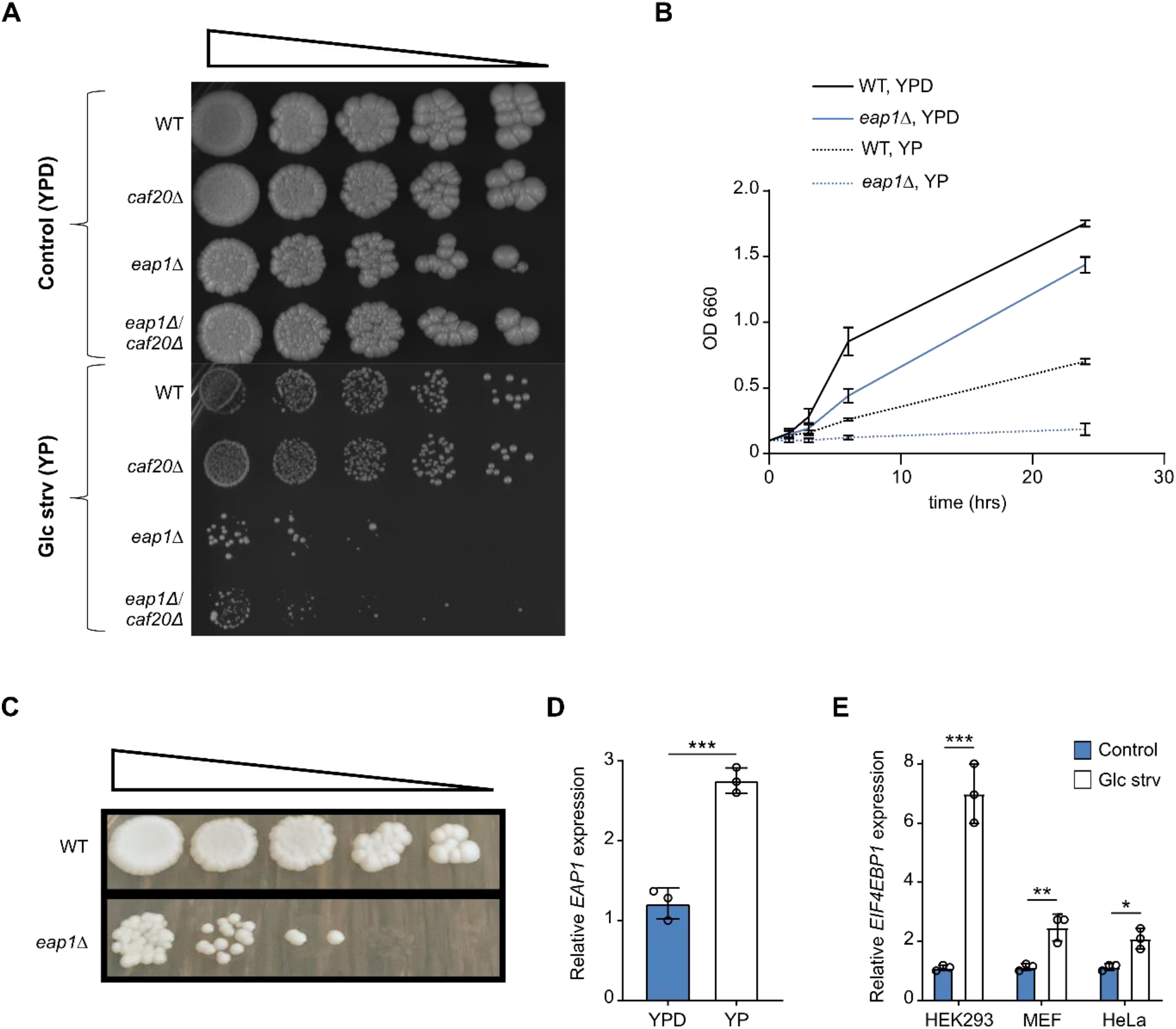
Yeast 4EBP orthologue EAP1 promotes survival of yeast cells under glucose deprivation. (A) WT, *eap1Δ, caf20Δ or eap1Δ*/*caf20Δ* strains were plated by serial dilution on solid complex medium with (YPD) or without (YP) 2% glucose at 37°C. (B) WT or *eap1Δ* were grown in liquid medium with (YPD) or without (YP) 2% glucose for the indicated times at which OD was measured at 30°C. (C) WT or *eap1Δ* were grown in liquid medium containing (YP) no glucose for 2 weeks at 30°C and were plated by serial dilutions onto complete YPD agar plates. (D) WT strains were grown in liquid medium with (YPD) or without (YP) 2% glucose at 30°C, and *EAP1* mRNA expression was analyzed by qRT-PCR. (E) HEK293, MEF and HeLa cells were grown in complete medium or glucose starved (Glc strv) for 24 hrs, and *EIF4EBP1* mRNA expression was analyzed by qRT-PCR. Where shown, data are reported as means ± SD with indicated significance (*p < 0.05, **p < 0.01, and ***p < 0.005).

Next, we determined whether *EAP1* and its orthologue *EIF4EBP1* respond to glucose deprivation, as part of a transcriptional program, as is expected from genes evolved to function during such stress. Indeed, we observed that levels of *EAP1* were induced following 24 hrs of glucose removal in yeast cultures (Fig. 5D) and that the expression of *EIF4EBP1* was similarly increased upon glucose starvation in three mammalian cell lines (Fig. 5E). Together, these data reveal that 4EBP1/2 are conserved components of the biological response to glucose starvation.

### 4EBP1/2 promote oncogenic transformation by mitigating oxidative stress and controlling ACC1 level

The cellular response to glucose starvation is closely linked to oncogenic transformation and tumorigenicity, as similar mechanisms governing redox balance and fatty acid synthesis are involved in these biological processes (Jeon et al., 2012; Schafer et al., 2009; Truitt et al., 2015). While 4EBP1/2 are required for oncogenic RAS transformation of primary fibroblasts (Petroulakis et al., 2009), it is not known whether 4EBP1/2 support transformation by other oncogenes or contribute to the maintenance of the oncogene-transformed state, as is expected if 4EBP1/2 indeed promote survival during energetic stress (Jeon et al., 2012). Using soft agar colony formation assays, we uncovered that 4EBP1/2 is necessary for HER2 transformation of mouse mammary epithelial cells (NT2197) in vitro (Fig. 6A). Similarly, 4EBP1/2 kd restricted the ability of KRAS^V12^-transformed, immortalized NIH 3T3 fibroblasts to form colonies in soft agar (Fig. S4A). Conversely, overexpression of 4EBP1^AA^ in HeLa cells led to a significant increase in colony formation in soft agar compared to control (EV) (Fig. 6B). Thus, these data demonstrate that 4EBP1/2 pro-tumorigenic functions are neither restricted to the *RAS* oncogene nor only to initiation of cellular transformation.

**Figure 6.**
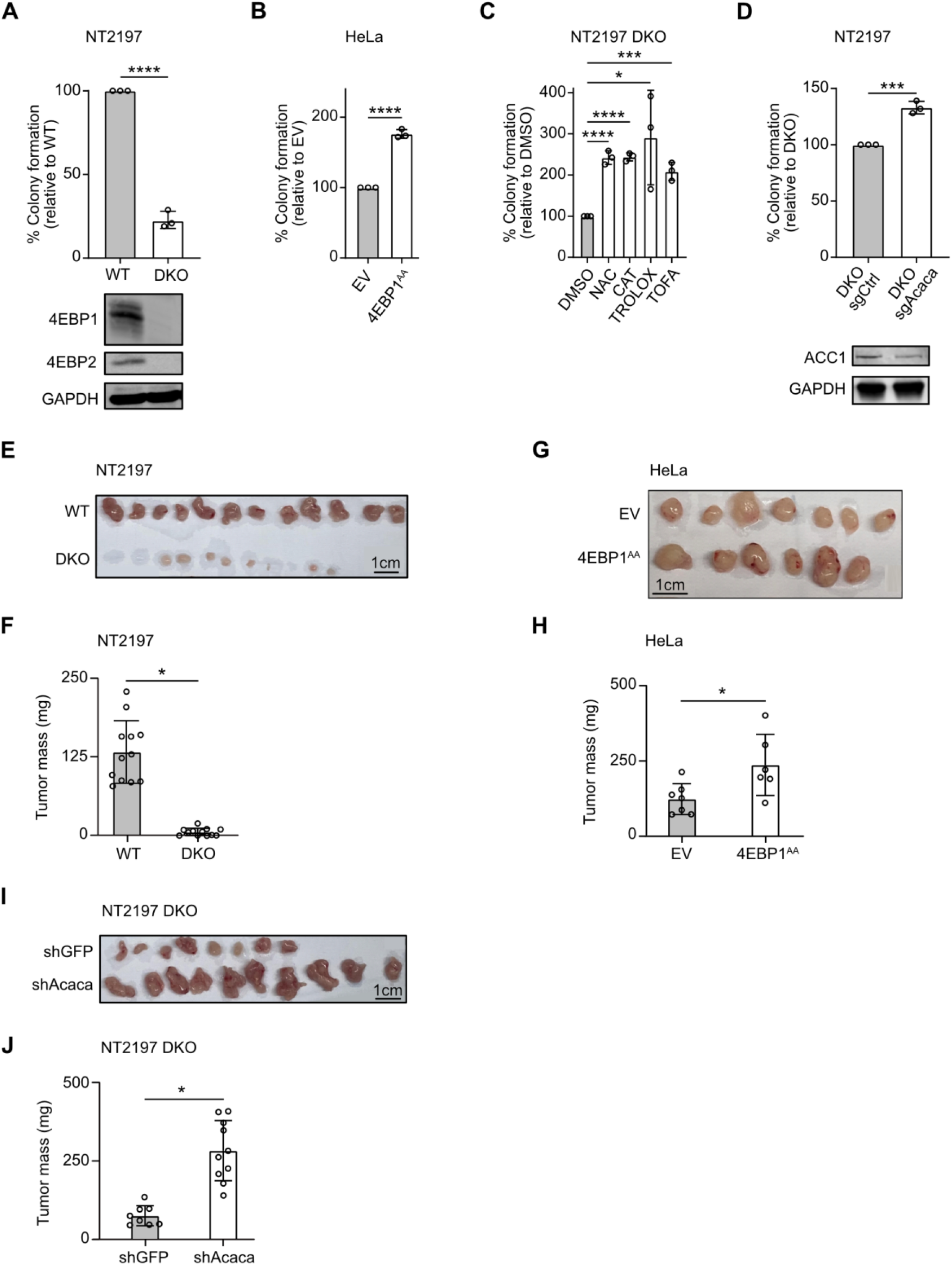
4EBP1 supports oncogenic transformation in vitro and in vivo. (A, B) WT and 4EBP1/4EBP2 DKO NT2197 (A), or empty vector (EV) and 4EBP1^AA^ expressing HeLa cells (B) were grown in soft agar for 21 days. Colonies and single cells were counted, and colony formation efficiency was calculated and normalized to respective control. Protein expression of 4EBP1 and 4EBP2 was analyzed by immunoblotting. (C) 4EBP1/4EBP2 DKO NT2197 cells were grown in soft agar for 21 days and treated with DMSO, NAC, CAT, Trolox or TOFA. Colonies and single cells were counted, and colony formation efficiency was calculated and normalized to DMSO. (D) Control (sgCtrl) and ACC1 targeting CRISPRi (sgAcaca) 4EBP1/4EBP2 DKO NT2197 cells were grown in soft agar for 21 days. Colony formation efficiency and proteins level were analyzed as in (A, B). (E, F) WT or 4EBP1/4EBP2 DKO NT2197 cells were injected in the mammary fat pad of NOD *SCID* gamma mice. Tumors were harvested, photographed (E) and weighed (F). (G, H) EV or 4EBP1^AA^ expressing HeLa cells were injected in the flank of NOD *SCID* gamma mice. Tumors were harvested, photographed (G) and weighed (H). (I, J) Control (shGFP) or stable ACC1 knock down (shAcaca) 4EBP1/4EBP2 DKO NT2197 cells were injected in the flank of NOD *SCID* gamma mice. Tumors were harvested, photographed (I) and weighed (J). Where shown, data are reported as means ± SD with indicated significance (*p < 0.05, ***p < 0.005, and ****p < 0.0001).

To determine how 4EBP1/2 deficiency inhibits oncogenic transformation, we assessed the possible involvement of oxidative stress and uncontrolled fatty acid synthesis. Treatment of 4EBP1/2 DKO NT2197 cells with antioxidants – CAT, NAC or Trolox – or with the ACC inhibitor TOFA rescued colony formation in soft agar (Fig. 6C), while it had no observable effect on 4EBP1/2 WT NT2197 cells (Fig. S4B). Similarly, antioxidants treatment restored colony formation of NIH 3T3 KRAS^V12^ 4EBP1/2 kd cells in soft agar (Fig. S4C). Importantly, we found that clustered regularly interspaced short palindromic repeats interference (CRISPRi)-mediated kd of Acc1 expression was sufficient to restore colony formation of 4EBP1/2 deficient NT2197 cells (Fig. 6D). Thus, we conclude that 4EBP1/2 support oncogenic transformation by negatively regulating ACC1 and oxidative stress.

We next tested our model in vivo and found that 4EBP1/2 DKO NT2197 cells were unable to form any palpable tumors when injected in the fat pad of immunocompromised mice, in sharp contrast to WT NT2197 cells, which formed sizeable tumors in all mice (12/12) (Fig. 6E&F). In line with this, overexpression of 4EBP1^AA^ in HeLa cells promoted tumor growth relative to control (EV) HeLa cells when injected in the flanks of immunocompromised mice (Fig. 6G&H). To ascertain the contribution of ACC1 to the observed phenotype in NT2197 tumors in vivo, we assessed the impact of targeting ACC1 expression on the growth of 4EBP1/2 DKO NT2197 tumors. Acc1 kd (shAcaca) in 4EBP1/2 DKO NT2197 cells led to a major increase of tumor mass as compared to control (shGFP 4EBP1/2 DKO NT2197) tumors (Fig. 6I&J). Collectively, these data support a model whereby 4EBP1/2 promote oncogenic transformation, tumorigenicity and survival during glucose starvation through a common mechanism entailing reduced ACC1 expression to restrain fatty acid synthesis and, thus, oxidative stress.

### 4EBP1 is clinically relevant and functional in brain tumors

Having found that 4EBP1/2 promote both survival upon glucose starvation, a condition commonly encountered in solid tumors, and oncogenic transformation, we further investigated the clinical relevance of 4EBP1/2 in cancer. It has been reported that *EIF4EBP1* is overexpressed in numerous tumor entities of TCGA and high *EIF4EBP1* expression in all TCGA tumors pooled together is associated with poor patient outcome (Wu and Wagner, 2021). Reanalysis of TCGA and GTEx datasets indicates that, together with our previous report (Hauffe et al., 2022), *EIF4EBP1* is overexpressed in 17 different tumor types compared to corresponding normal tissues (Fig. S5A). In contrast, *EIF4EBP2* is only overexpressed in 3 out of these 17 tumor types in TCGA datasets (Fig. S5B), leading us to focus our analyses on *EIF4EBP1*. Indeed, we uncovered that high *EIF4EBP1* expression correlated with significantly decreased overall survival in three different tumor types (Fig. S5C-E), including glioma, highlighting *EIF4EBP1* expression as a potential prognostic biomarker in these tumor entities.

Glucose levels are low in the interstitial compartment of the brain as compared with blood (Fellows and Boutelle, 1993; Gruetter et al., 1992). To survive in the pre-existing low glucose microenvironment of the brain, glioma tumor cells or cancer cells that metastasize to the brain acquire resistance to glucose starvation and/or use alternative energy sources (Chen et al., 2015; Flavahan et al., 2013). To further investigate the relevance of *EIF4EBP1* in cancer, we thus turned our attention towards malignant gliomas, the most common form of brain tumor, which is characterized by glucose deprivation (Tanaka et al., 2021). *EIF4EBP1* expression is increased with gliomas tumor grade, with the highest expression being found in the most aggressive grade, CNS World Health Organisation (WHO) grade 4 glioblastoma [relative to lower grade gliomas (CNS WHO grades 2 and 3)] (Fig. 7A). Furthermore, high *EIF4EBP1* expression was associated with reduced overall survival in one additional independent and non-overlapping glioma cohort (Fig. 7B). This trend was similarly observed in glioblastoma patients (Fig. 7C), illustrating the clinical relevance of *EIF4EBP1* in such a highly malignant form of adult brain tumor. Additionally, we analyzed proteomic data obtained from glioblastoma patient samples and observed that 4EBP1 protein is overexpressed in glioblastoma tissues compared to non-tumorigenic brain tissues (NTBT) (Fig. S5F). Importantly, 4EBP1 protein levels were found to be negatively correlated with ACC1 protein expression in glioblastoma (Fig. 7D), supporting our model whereby 4EBP1 represses ACC1 synthesis.

**Figure 7.**
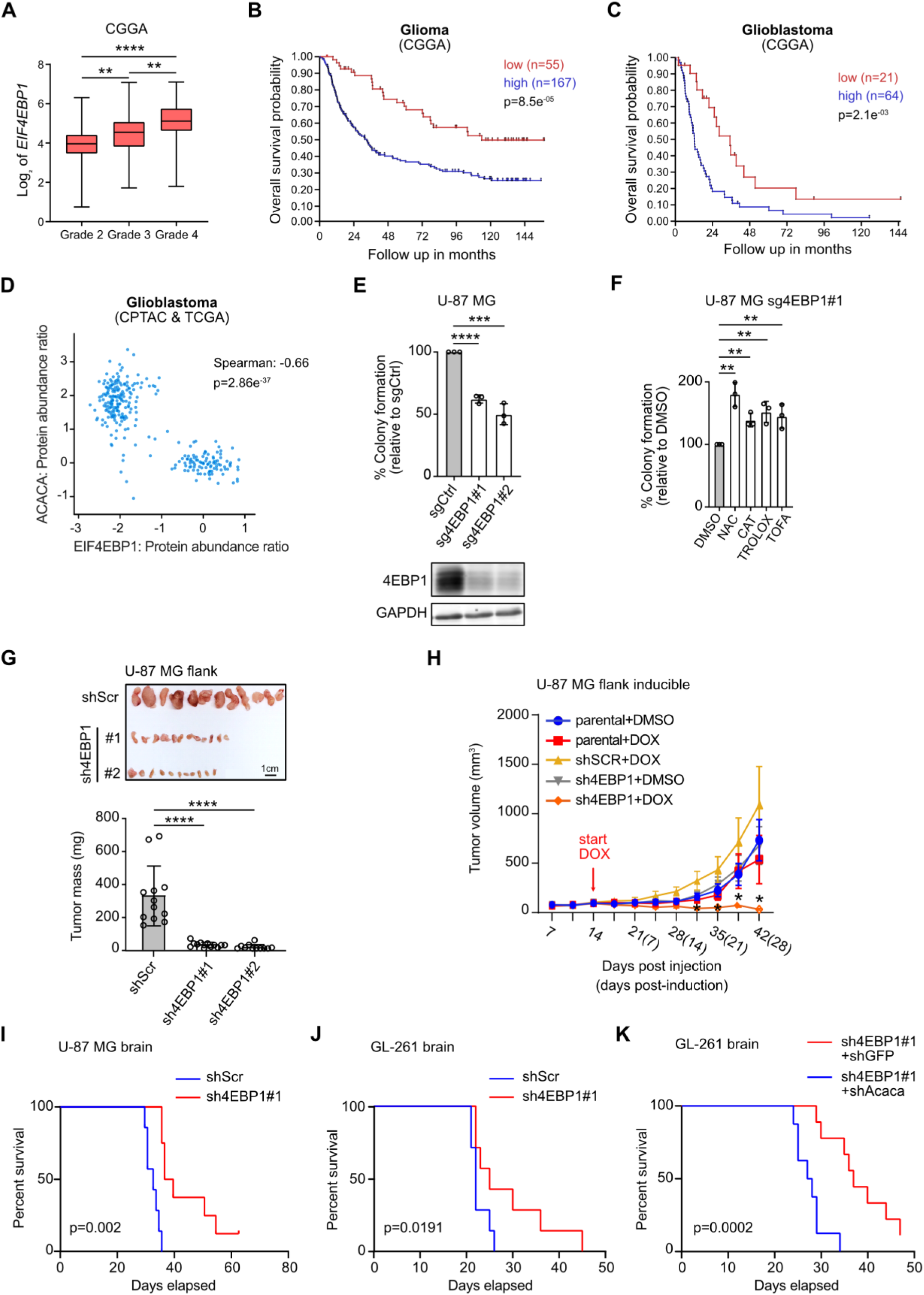
4EBP1 has clinical relevance in glioma and promotes glioma tumorigenesis. (A) Expression levels of *EIF4EBP1* per glioma grade in the CGGA cohort. p values were calculated using an unpaired and two-tailed parametric *t* test. (B, C) Kaplan-Meier survival estimates of overall survival of glioma (B) or glioblastoma (C) patients stratified by their *EIF4EBP1* mRNA levels (cut off first quartile) in the CGGA cohort. p values were calculated using a log rank test. (D) Expression levels of 4EBP1 protein in glioblastoma patient samples plotted against the expression levels of ACC1 protein using CPTAC and TCGA GBM proteomic data. Co-expression levels were quantified by calculating the Spearman’s rank correlation coefficient. (E) Control (sgCtrl) or *EIF4EBP1* targeting CRISPRi (sg4EBP1#1 and #2) U-87 MG cells were grown in soft agar for 21 days. Colonies and single cells were counted, and colony formation efficiency was calculated and normalized to control. Data are reported as means ± SD with indicated significance. (F) Sg4EBP1#1 U-87 MG cells were grown in soft agar for 21 days and treated with DMSO, NAC, CAT, Trolox or TOFA. Colonies and single cells were counted, and colony formation efficiency was calculated and normalized to DMSO. Data are reported as means ± SD with indicated significance. (G) Control (shScr) or stable 4EBP1 knock down (sh4EBP1#1 and #2) U-87 MG cells were injected in the flank of NOD *SCID* gamma mice. Tumors were harvested, photographed and weighed. (H) ShScr or stable inducible 4EBP1 knock down (sh4EBP1) U-87 MG cells were injected in the flank of NOD *SCID* gamma mice. When tumors reached 100 mm^3^, mice were given doxycycline (DOX) or vehicle. Tumor volumes were measured at the indicated times. (I) ShScr (n=7) or sh4EBP1#1 (n=8) U-87 MG cells were injected intracranially in NOD *SCID* gamma mice. Survival of mice was monitored post injection. p value was calculated using a log rank test. (J) ShScr (n=10) or sh4EBP1#1 (n=10) GL-261 cells were injected intracranially in C57WT mice. Survival of mice was monitored post injection. p value was calculated using a log rank test. (K) Sh4EBP1#1 containing shGFP (n=9) or shAcaca (n=8) GL-261 cells were injected intracranially in C57WT mice. Survival of mice was monitored post injection. p value was calculated using a log rank test. Indicated significance are **p < 0.01, and ***p < 0.005, and ****p < 0.0001.

To functionally dissect the role of 4EBP1 in glioma, we analyzed the impact of 4EBP1 kd on the tumorigenic potential of human and mouse glioma cells, U-87 MG and GL-261, respectively. We first confirmed that 4EBP1 kd sensitizes such glioma cells to glucose starvation-induced cell death (Fig. S5G&H), and severely restricted the ability of glioma cells to form colonies in soft agar (Fig. 7E&S6A) as with the other previously demonstrated cell lines. Importantly, inhibiting ACC with TOFA or supplementing cells with antioxidants rescued colony formation of 4EBP1 deficient glioma cells (Fig. 7F, S6B-D), which was not observed with corresponding 4EBP1 proficient cells (Fig. S6E&F). These data indicate that in glioma cells, 4EBP1 promotes tumorigenicity in vitro by means of controlling redox balance and fatty acid synthesis.

We next evaluated pro-tumorigenic functions of 4EBP1 in glioma cells in vivo by first injecting control (shScr) and 4EBP1 kd U-87 MG cells to the flanks of NOD *SCID* mice. 4EBP1 depleted cells grew to significantly smaller tumors as compared to controls (Fig. 7G). To determine whether 4EBP1 is also important for tumor maintenance, we used a doxycycline-inducible shRNA system to target 4EBP1 expression in established tumors. Namely, engineered inducible 4EBP1 kd U-87 MG models were injected into the flanks of NOD *SCID* mice, and once tumor size reached 100 mm^2^, 4EBP1 kd was induced by adding doxycycline to their drinking water. We observed an inhibition of tumor growth in doxycycline treated mice harboring the sh4EBP1 U-87 MG tumors but not in shScr U-87 MG tumors or tumors in mice unexposed to doxycycline (Fig. 7H). These data suggest that 4EBP1 promotes growth of established glioma tumors in vivo. To further corroborate our findings that 4EBP1 supports glioma tumor growth, specifically when localized in the brain, we performed orthotopic injection of control and sh4EBP1 U-87 MG cells. While both control and 4EBP1 deficient cells generated tumors, mice bearing sh4EBP1 U-87 MG tumors survived longer as compared with controls (Fig. 7I).

Finally, to assess 4EBP1 function in an immunocompetent mouse model, we injected control and sh4EBP1 GL-261 models in brains of C57WT mice. Mice carrying sh4EBP1 GL-261 tumors showed a significant extension of survival compared to mice with control (shScr) tumors (Fig. 7J), suggesting that 4EBP1 promotes glioma aggressiveness even in the presence of a functional immune system. Importantly, Acc1 kd enhanced tumor aggressiveness of orthotopically injected sh4EBP1 GL-261 cells, as evidenced by reduced mice survival (Fig. 7K). This supports the model whereby 4EBP1 function is exploited by glioma cells to reduce Acc1 expression to promote tumor aggressiveness. Collectively, our data highlight that *EIF4EBP1* is linked to malignancy and poor outcome in various human tumor types, including glioma, and that 4EBP1 exerts a pro-tumorigenic function in glioblastomas by reducing ACC1 expression.

## DISCUSSION

### 4EBP1/2 are evolutionarily conserved factors promoting cell survival under glucose deprivation

Glucose starvation represents a physiological stress that necessitates proper cellular responses to prevent cell death. Here we report that 4EBP1/2 promote cell survival during glucose starvation, and demonstrate that this biological function is conserved in human, mouse and yeast cells. This is in line with the proposed role of 4EBP in facilitating *Drosophila melanogaster* viability under dietary restriction (Teleman et al., 2005; Tettweiler et al., 2005; Zid et al., 2009). While the main known functions of 4EBP1/2 are to restrict cell proliferation and overall protein synthesis (Dowling et al., 2010; Sonenberg and Hinnebusch, 2009), we found that this is not the case under glucose starvation. Conversely, our data support that 4EBP1/2 act as pro-survival factors under glucose deprivation, but not under amino acid starvation, serum starvation or pharmacological inhibition of mTORC1 (Dowling et al., 2010), highlighting that the type of stress encountered by cells may dictate 4EBP1/2 cellular functions. It is worth noting the 4EBPs yeast ortholog Eap1p shows no sequence homology with mammalian 4EBPs, apart from a conserved YXXXXL eIF4E binding motif, thus indicating that 4EBP1/2 pro-survival function is conserved despite a lack of sequence conservation.

We found that *EIF4EBP1* has evolved as a glucose starvation-responsive gene. Both yeast *EAP1* and mammalian *EIF4EBP1* expression are induced by glucose starvation, in line with observations made in glucose-starved adipocytes (Agudelo et al., 2021) and in muscle tissue of food-deprived mice (Jagoe et al., 2002). The findings that glucose starvation induced expression of another pro-survival factor and negative regulator of mRNA translation, namely eEF2K (Leprivier et al., 2013), are supportive of a model whereby negative regulators of mRNA translation, and 4EBP1/2 in particular, have evolved to protect cells against glucose starvation.

### 4EBP1/2 act as metabolic switches by translationally restricting fatty acid synthesis

The cellular response to glucose starvation encompasses profound metabolic reprogramming, during which anabolic processes are blocked and catabolic processes are activated (Caro-Maldonado, 2011). Notably, our findings uncovered that 4EBP1/2 are key mediators of the metabolic switch induced by glucose withdrawal by restricting fatty acid synthesis activity. This allows cells to preserve NADPH levels and maintain the redox balance, similar to the reported function of the energy sensor AMPK (Jeon et al., 2012). However, unlike AMPK which regulates fatty acid synthesis at the posttranslational level, i.e. by phosphorylating and inhibiting the fatty acid synthesis rate limiting enzyme ACC1, we found that 4EBP1/2 control this process at the translational level, highlighting 4EBP1/2 as previously unrecognized translational regulators that fine tune redox balance according to the intracellular energy state. The ability of 4EBP1/2 to modulate metabolic processes, such as proliferation and mitochondrial activity, relies on their ability to selectively restrict the translation of specific transcripts. These include pro-proliferative *cyclin D3* and *ornithine decarboxylase* (Dowling et al., 2010) as well as *ATP5O* and *TFAM* which encode mitochondrial proteins (Morita et al., 2013). In addition to these mechanisms, we uncovered that in response to glucose starvation, 4EBP1/2 restrain fatty acid synthesis by selectively inhibiting the synthesis of ACC1. It was previously reported that eIF4E selectively promotes *ACACA* translation in T cells (Ricciardi et al., 2018) and in liver tissue of mice fed with a high-fat diet (Conn et al., 2021). In particular, the transition of CD4+ T cells from quiescence to activation, which metabolically mirror changes in glucose availability, is driven by eIF4E-promoted *ACACA* translation which is dependent upon a 5’UTR of *ACACA* (Ricciardi et al., 2018). Similarly, we report that a second 5’UTR of *ACACA* supports 4EBP1/2-mediated control of *ACACA* translation, highlighting that this *ACACA* 5’UTR represents a genetic element linking fatty acid synthesis activity to the energetic state of the cell. Taken together, the regulation of cell metabolism by 4EBP1/2, through inhibition of fatty acid synthesis, proliferation (Dowling et al., 2010) and mitochondrial activity (Morita et al., 2013), is compatible with 4EBP1/2 acting as stress-responsive metabolic switches that steer cells towards a more quiescent or low energy state.

Altogether, our data support a model whereby in addition to AMPK-mediated phosphorylation of ACC1, cells evolved a parallel mechanism to inhibit ACC1 in response to energetic stress through 4EBP1/2-mediated translational repression of *ACACA* mRNA. These may represent two distinct but complementary mechanisms, given that phosphorylation occurs rapidly and is easily reversible by the action of phosphatases, while repression of mRNA translation occurs more slowly but may be a more durable and efficient means for blocking ACC1. Notably, in bacteria, ACC is also regulated at both the translational and post-translational levels in response to glucose availability (Broussard et al., 2013), in agreement with the possibility that both modes of ACC1 regulation are evolutionarily conserved.

### 4EBP1 exerts a pro-tumorigenic function in glioma

The role of 4EBP1 in cancer remains unclear (Musa et al., 2016). While 4EBP1 exhibits a tumor suppressive function in mouse models of lymphoma, head and neck squamous cell carcinoma and prostate cancer (Ding et al., 2018; Wang et al., 2019), 4EBP1 KO mice do not develop tumors per se, thus excluding 4EBP1 as a bona fide tumor suppressor (Tsukiyama-Kohara et al., 2001). Furthermore, 4EBP1 has been shown to exert pro-tumorigenic functions, as it is required for oncogenic RAS transformation (Petroulakis et al., 2009), promotes breast cancer development in vivo (Braunstein et al., 2007), and its expression correlates with poor patient outcome in several tumor types at the transcriptional level (Wu and Wagner, 2021). Here, our data further support a pro-tumorigenic function of 4EBP1, as we uncovered that 4EBP1 mediates HER2 transformation of mouse mammary epithelial cells and tumorigenicity of glioma cells in vitro and in vivo.

The role of 4EBP1 in cancer is likely determined by the levels of metabolic stress present in tumors, such that 4EBP1 acts as a pro-tumorigenic factor within metabolically challenged tumor environments, as was previously proposed for AMPK (Chhipa et al., 2018; Eichner et al., 2019; Faubert et al., 2014). In particular, glucose concentrations in the brain are low compared to plasma (Fellows and Boutelle, 1993; Gruetter et al., 1992), and glioblastoma are characterized by further reduction of glucose levels in the central region of the tumor (Tanaka et al., 2021). For glioma tumor cells or breast cancer cells that metastasize to the brain, a precondition for survival under such a low glucose microenvironment is the acquisition of resistance to glucose starvation (Chen et al., 2015; Flavahan et al., 2013), which may adversely select for more highly tumorigenic cancer cell clones (Flavahan et al., 2013). With this perspective, and based on our data, we propose that 4EBP1 confers glioma cells the ability to adapt to such metabolic stress by preserving cellular redox balance and restricting ACC1 expression, by co-opting the mechanisms of 4EBP1 function in response to glucose deprivation. This is in line with the proposed function of AMPK in mediating cell survival under glucose starvation and tumorigenesis through inhibition of ACC1 and prevention of oxidative stress (Jeon et al., 2012). Therefore, 4EBP1 represents a metabolic regulator exploited by cancer cells to adapt to the adverse conditions of the tumor microenvironment.

It is worth noting that since 4EBP1 is post-translationally inhibited by mTORC1, which is overactive in numerous cancers, it is assumed that 4EBP1 is inactive in tumors as evidenced by increased levels of phosphorylated 4EBP1 reported in various tumor tissues (Musa et al., 2016). However, the amount of total 4EBP1 protein, which is a contributing factor, is rarely monitored. Importantly, the activity of 4EBP1 in glioblastoma was shown to be directly dependent on the proximity to blood vessels, with highest 4EBP1 activity detected in areas furthest from blood vessels, corresponding to oxygen and glucose deprived areas (Kumar et al., 2019). This raises the possibility that upregulation of *EIF4EBP1*, as observed in numerous cancer types (Wu and Wagner, 2021), leads to increased 4EBP1 activity in metabolically challenged tumor areas. It is also worth noting that oncogenic transcription factors, such as ETS1, MYBL2, MYC and MYCN (Hauffe et al., 2022; Tameire et al., 2019; Voeltzke et al., 2022) promote *EIF4EBP1* overexpression, further supporting the clinical relevance of *EIF4EBP1* as a pro-tumorigenic gene. Together, our findings suggest that 4EBP1 may represent a therapeutic target in metabolically challenged tumor types, while warranting caution on the use of mTOR inhibitors in these cancers. To summarize, our findings reveal that 4EBP1/2 inhibit fatty acid synthesis during glucose starvation and that this particular function is exploited by tumor cells for their own selective advantage.

## ACKNOWLEDGMENTS

We would like to thank Maya Bar and Björn Stork for helpful discussions. We would like to thank Marc B. Hershenson (University of Michigan) for providing us the MSCV puro-4EBP1 (T37A/T46A) plasmid. B.R. was supported by the Israel Cancer Association (grant No. 20220143), the ISRAEL SCIENCE FOUNDATION (grant No. 1436/19) and the NIBN. G.L. was supported by grants from the Deutsche Forschungsgemeinschaft (grant no. LE 3751/2-1), the German Cancer Aid (grant no. 70112624), the Elterninitiative Düsseldorf e.V. (grant no. 701910003), the Dr. Rolf M. Schwiete Stiftung (grant no. 2020-018) and the Research Commission of the Medical Faculty, Heinrich Heine University Düsseldorf (grant no. 2016-056 and 2020-044). LH was funded by the Dr. Rolf M. Schwiete Stiftung (grant no. 2020-018). The laboratory by T.G.P.G. is supported by grants from the by the Matthias-Lackas Foundation, the Dr. Leopold and Carmen Ellinger Foundation, the German Cancer Aid (DKH-70112257 and DKH-70114111), the SMARCB1 association, the Boehringer-Ingelheim Foundation, the Deutsche Forschungsgemeinschaft (DFG-458891500), the Ministry of Science and Education (BMBF MCORESYS-014 SMART-CARE (031L0212B)), and the Barbara und Wilfried Mohr Foundation. C.M.F. received a scholarship from the German Cancer Aid. S.-M.F. acknowledges funding from the European Research Council under the ERC Consolidator Grant Agreement n. 771486–MetaRegulation, FWO – Research Projects, KU Leuven – FTBO, King Baudouin Foundation, Beug Foundation and Fonds Baillet Latour. The results published here are in part based on data generated by TCGA Research Network (https://www.cancer.gov/tcga).

## AUTHOR CONTRIBUTIONS

Conceptualization, T.L., K.A., K.V., L.H., B.R., and G.L.; Methodology, T.L., K.A., K.V., L.H., K.S., B.H., and G.L.; Investigation, T.L., K.V., L.H., K.A., M.S., R.M., K.S., M.P., K.Vr., S.C., C.M.F., C.H., A.K., B.H., K.B., D.P., Z.B., B.R., and G.L.; Resources, A.V-T., U.K., and M.E.; Writing – Original Draft, B.R. and G.L.; Writing – Review & Editing, K.V., L.H., K.A., T.L., S-M.F., B.R., J.K.M.L. and G.L.; Visualization, L.H., K.V., B.R. and G.L.; Supervision, M.R., T.G.P.G., A.S.R., S-M.F., A.S., G.R., B.R. and G.L.; Funding Acquisition, B.R., J.K.M.L. and G.L.

## SUPPLEMENTAL INFORMATION

The Supplemental Information includes six figures, Materials and Methods.

