## Supplementary material for "4EBP1/2 support tumorigenicity and cell survival during energetic stress by translationally regulating fatty acid synthesis": SUP figs and methods

- Supplementary figures
- Supplementary tables with sequences, materials and antibodies
- Experimental model and subject details
- Methods details
- References

Suppl Figure 1

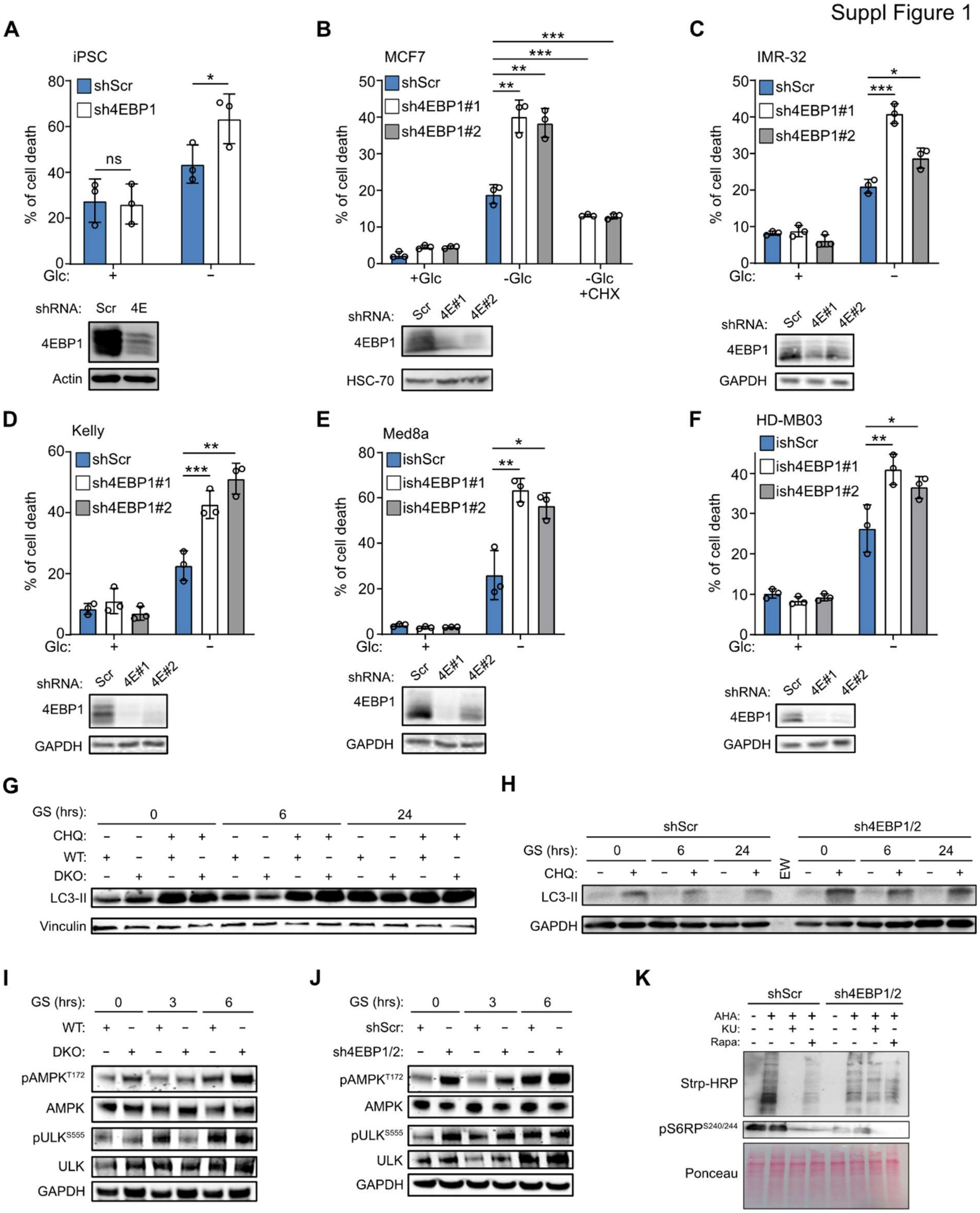

**Supplementary Figure 1. Related to Figure 1.**

(A-F) Control (shScr) and stable 4EBP1 knock down (sh4EBP1) iPSC (A), shScr, sh4EBP1#1 and #2 MCF7 (B), IMR-32 (C), and Kelly (D), or inducible control (ishScr) and stable 4EBP1 knock down (ish4EBP1) Med8a (E) and HD-MB03 cells (F) were grown in complete medium or glucose (Glc) starved for 48 hrs. Med8a and HD-MB03 cells were treated with 1 μg/ml doxycycline for 72 hrs. MCF7 cells were treated or not with cycloheximide (CHX) for 48 hrs. Cell death was measured by PI staining and flow cytometry.

(G, H) WT and 4EBP1/4EBP2 DKO MEF (G), or shScr and sh4EBP1/2 HEK293 cells (H) were grown in complete medium or glucose starved (GS) for the indicated times with or without chloroquine (CHQ). Cell lysates were analyzed by immunoblotting using antibodies against the indicated proteins.

(I, J) WT and 4EBP1/4EBP2 DKO MEF (I), or shScr and sh4EBP1/2 HEK293 cells (J) were grown in complete medium or glucose starved (GS) for the indicated times. Cell lysates were analyzed by immunoblotting using antibodies against the indicated proteins.
(K) ShScr and Sh4EBP1/2 HEK293 cells were grown in complete media, treated or not with rapamycin (Rapa) or Ku-0063794 (KU), and labeled with azidohomoalanine (AHA). Levels of AHA-labelled proteins were detected by immunoblotting with a streptavidin conjugate.
Where shown, data are reported as means ± SD with indicated significance (*p < 0.05, **p < 0.01, and ***p < 0.005).

Suppl Figure 2

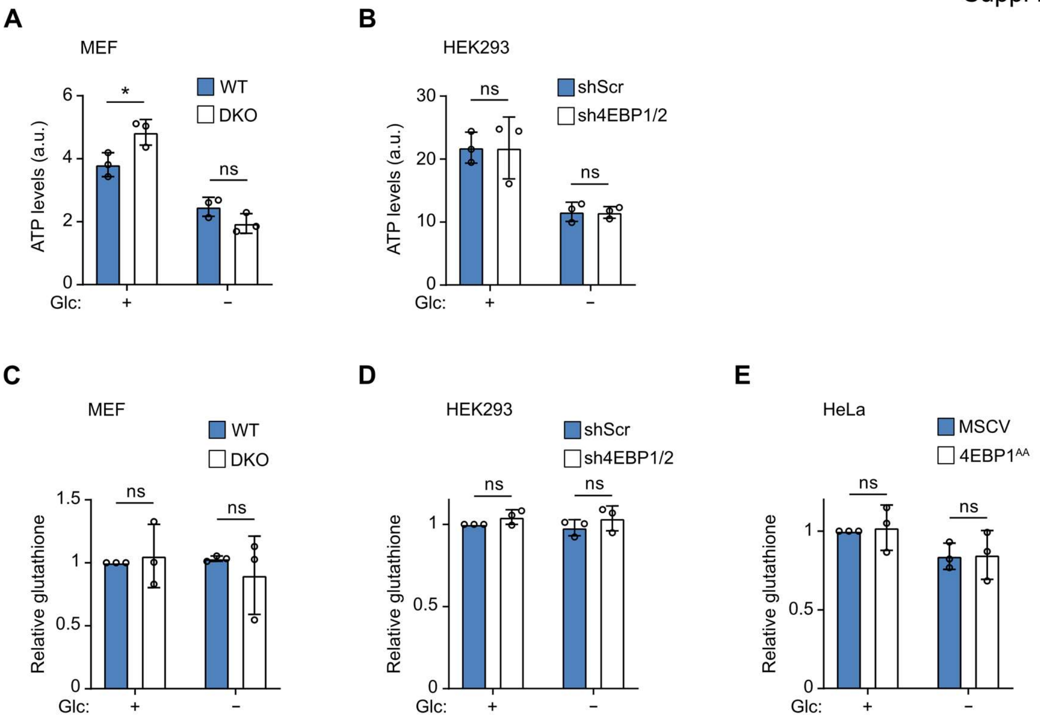

**Supplementary Figure 2. Related to Figure 2.**
(A, B) WT and 4EBP1/4EBP2 DKO MEF (A), or shScr and sh4EBP1/2 HEK293 cells (B) were grown in complete medium or glucose (Glc) starved for 24 hrs, and ATP levels were measured by GC-MS. A.u.: arbitrary units.
(C-E) WT and 4EBP1/4EBP2 DKO MEF (C), shScr and sh4EBP1/2 HEK293 cells (D), or MSCV or 4EBP1AA overexpressing HeLa cells (E) were grown in complete medium or glucose (Glc) starved for 24 hrs, and total glutathione was measured.
Where shown, data are reported as means ± SD with indicated significance (*p < 0.05).

Suppl Figure 3

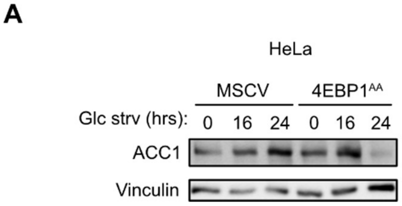

**Supplementary Figure 3. Related to Figure 4.**
(A) MSCV or 4EBP1AA overexpressing HeLa cells were grown in complete medium or glucose starved (Glc strv) for the indicated times, and analyzed by immunoblotting using antibodies against ACC1 and Vinculin as reference.

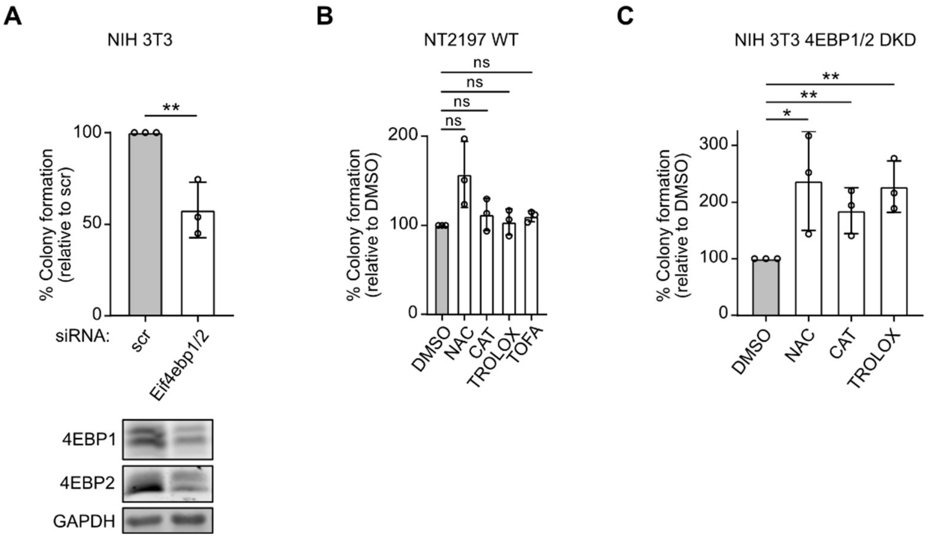

**Supplementary Figure 4. Related to Figure 6.**

(A) NIH 3T3 transfected with control (scr), or *Eif4ebp1* and *Eif4ebp2* targeting siRNAs (Eif4ebp1/2) were grown in soft agar for 21 days. Colonies and single cells were counted, and colony formation efficiency was calculated.

(B) 4EBP1/2 WT NT2197 cells were grown in soft agar for 21 days and treated with DMSO, NAC, CAT, Trolox or TOFA. Colonies and single cells were counted, and colony formation efficiency was calculated.

(C) NIH 3T3 transfected with *Eif4ebp1* and *Eif4ebp2* targeting siRNAs (4EBP1/2 DKD) were grown in soft agar for 21 days and treated with DMSO, NAC, CAT or Trolox. Colonies and single cells were counted, and colony formation efficiency was calculated.

Where shown, data are reported as means ± SD with indicated significance (*$p < 0.05$, and **$p < 0.01$).

Suppl Figure 5

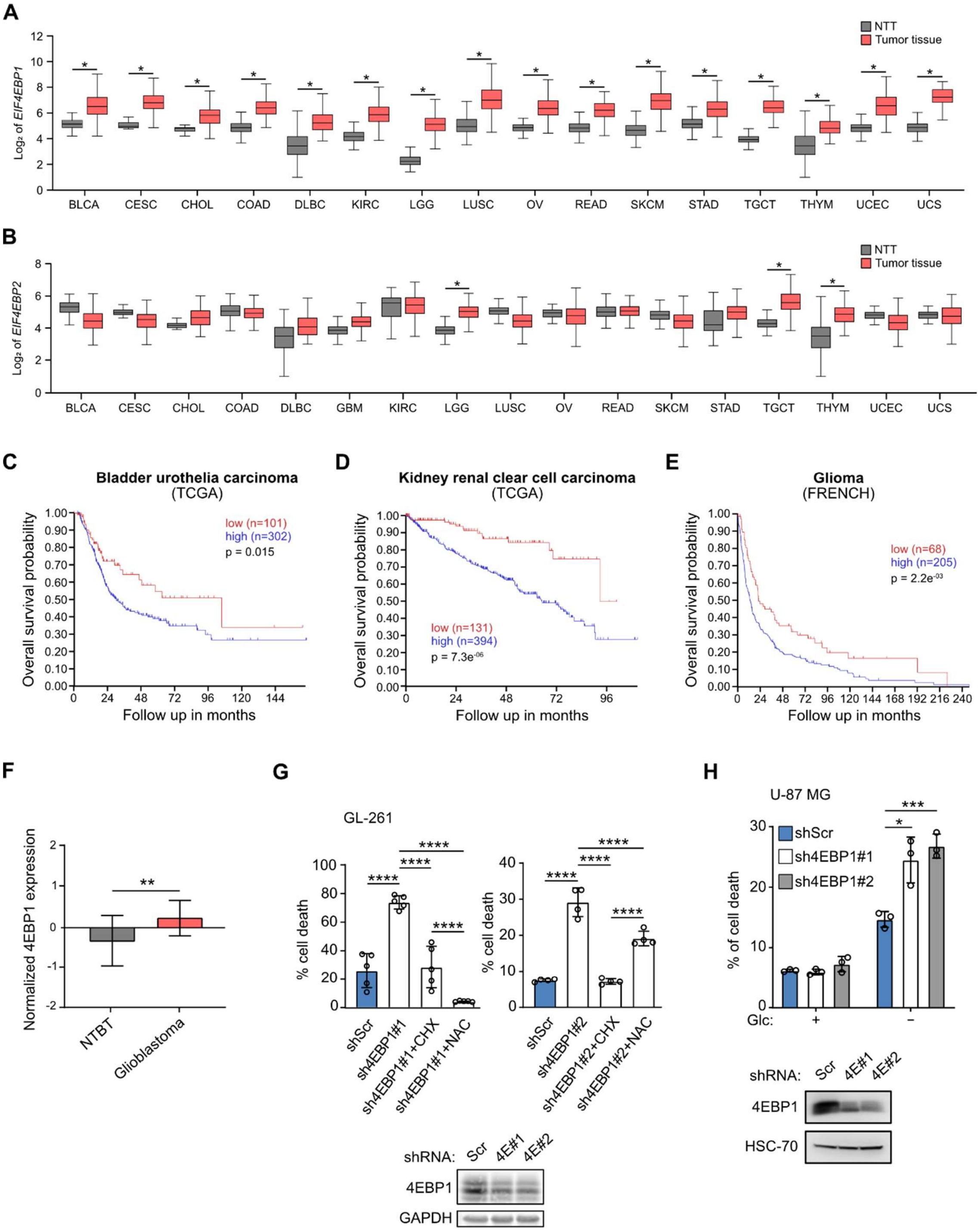

**Supplementary Figure 5. Related to Figure 7.**

(A, B) Expression levels of *EIF4EBP1* (A) and *EIF4EBP2* (B) in the indicated tumor entities and corresponding non-tumorigenic tissues (NTT) from TCGA. p values were calculated using an unpaired and two-tailed parametric *t* test. BLCA: bladder urothelial carcinoma, CESC: cervical squamous cell carcinoma and endocervical adenocarcinoma, CHOL: cholangiocarcinoma, COAD: colon adenocarcinoma, DLBC: lymphoid neoplasm diffuse large B-cell lymphoma, KIRC: kidney renal clear cell carcinoma, LGG: brain lower grade glioma, LUSC: lung squamous cell carcinoma, OV: ovarian serous cystadenocarcinoma, READ: rectum adenocarcinoma, SKCM: skin cutaneous melanoma, STAD: stomach adenocarcinoma, TGCT: testicular germ cell tumors, THYM: thymoma, UCEC: uterine corpus endometrial carcinoma, UCS: uterine carcinosarcoma.

(C, D, E) Kaplan-Meier survival estimates of overall survival of bladder urothelial carcinoma (C), kidney renal clear cell carcinoma (D), and glioma (E) patients stratified by their *EIF4EBP1* mRNA levels (cut off first quartile) in the indicated cohorts. p values were calculated using a log rank test.
(F) Expression levels of 4EBP1 protein in non-tumorigenic brain tissue (NTBT) and glioblastoma tissues from CPTAC GBM proteomic data. p value was calculated using a two-tailed Mann Whitney test.
(G) Control (shScr) and stable 4EBP1 knock down (sh4EBP1#1 and #2) GL-261 cells were grown in complete medium or glucose (Glc) starved with or without CHX or NAC for 48 hrs. Cell death was measured by PI staining and flow cytometry. The levels of the indicated proteins were analyzed by immunoblotting.
(H) ShScr and stable 4EBP1 knock down (sh4EBP1#1 and #2) U-87 MG cells were grown in complete medium or glucose (Glc) starved for 48 hrs. Cell death and protein levels were analyzed as in (G).
Where shown, data are reported as means ± SD with indicated significance (*p < 0.05, **p < 0.01, ***p < 0.005, and ****p < 0.0001).

Suppl Figure 6

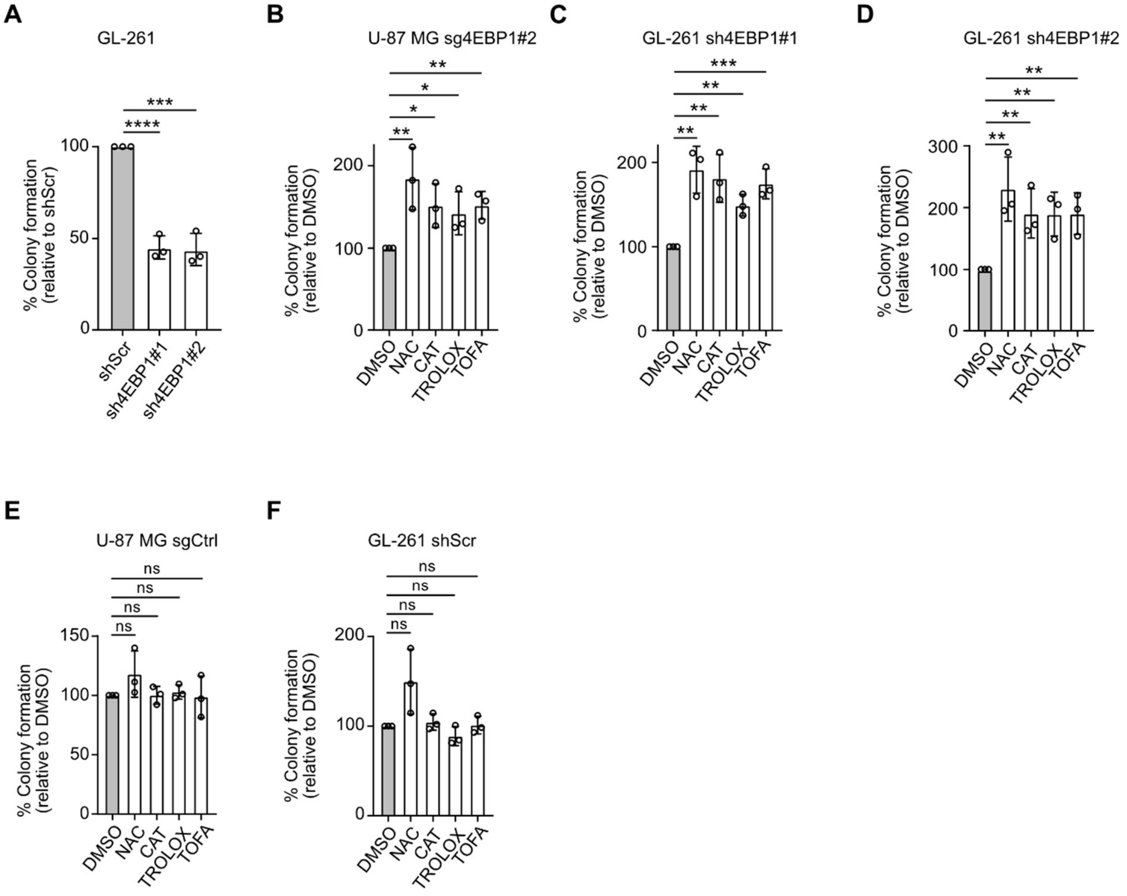

**Supplementary Figure 6. Related to Figure 7.**
(A) Control (shScr) and stable 4EBP1 knock down (sh4EBP1#1 and #2) GL-261 cells were grown in soft agar for 21 days. Colonies and single cells were counted, and colony formation efficiency was calculated and normalized to control.
(B-F) Sg4EBP1#2 U-87 MG (B), sh4EBP1#1 (C) and #2 (D) GL-261, sgCtrl U-87 MG (E) and shScr GL-261 cells (F) were grown in soft agar for 21 days and treated with DMSO, NAC, CAT, Trolox or TOFA. Colonies and single cells were counted, and colony formation efficiency was calculated and normalized to DMSO.
Where shown, data are reported as means ± SD with indicated significance (*p < 0.05, **p < 0.01, ***p < 0.005, and ****p < 0.0001).

**Supplementary table 1**: List of siRNA sequences

| siRNA label and target gene | siRNA sequence |
|---|---|
| Dharmacon – Horizon Discovery | |
| siGENOME Non-targeting Pool#1 | 5'- UGGUUUACAUGUCGACUAA -3' |
| | 5'- UAAGGCUAUGAAGAGAUAC -3' |
| | 5'- AUGUAUUGGCCUGUAUUAG -3' |
| | 5'- AUGAACGUGAAUUGCUCAA -3' |
| siGENOME mouse *Eif4ebp1* Smart Pool | 5'- GAACCAGGAUUAUCUAUGA -3' |
| | 5'- CAAAGGACCUGCCAGCCAU -3' |
| | 5'- GCGAUGAGCCUCCCAUGCA -3' |
| | 5'- CCAGCAGCCCGGAAGAUAA -3' |
| siGENOME mouse *Eif4ebp2* Smart Pool | 5'- GGGAGGAACACGAAUCAUU -3' |
| | 5'- UGAACAAUCAUGACAGGAA -3' |
| | 5'- GCACCGUGGCUAUCAGCGA -3' |
| | 5'- GUUGGACCGUCGCAAUUCU -3' |
| siGENOME human and mouse *EIF4E* siRNA#1 | 5'- CAUAUCCAGUUGUCUAGUA -3' |
| siGENOME human and mouse *EIF4E* siRNA#2 | 5'- GUGAUAAGAUAGCAAUAUG -3' |
| siGENOME human *ACACA* siRNA#1 | 5'- CAGCAAACCUGGAUUCUGA -3' |
| siGENOME human *ACACA* siRNA#2 | 5'- GCAAUUAGAUUCGUUGUCA -3' |
| siGENOME mouse *Acaca* siRNA#1 | 5'- AGAUAGAAUCAUCGAGUUU -3' |
| siGENOME mouse *Acaca* siRNA#2 | 5'- GGAUCAAGGAUAUUCGUAU -3' |
| ON-TARGETplus human *FASN* siRNA#1 | 5'- GAAGCACAUUGGCAAAGUC -3' |
| ON-TARGETplus human *FASN* siRNA#2 | 5'- CUUCCGAGAUUCCAUCCUA -3' |
| Sigma Aldrich | |
| mouse *Fasn* siRNA#1, sense | 5'- GUCAGAUCCUGGAACGAGA[dT][dT] -3' |
| mouse *Fasn* siRNA#1, antisense | 5'- UCUCGUUCCAGGAUCUGAC[dT][dT] -3' |
| mouse *Fasn* siRNA#2, sense | 5'- GUAAUGCUGGCCAAACUAA[dT][dT] -3' |
| mouse *Fasn* siRNA#2, antisense | 5'- UUAGUUUGGCCAGCAUUAC[dT][dT] -3' |

**Supplementary table 2**: List of CRISPRi sgRNA sequences

| sgRNA label and target gene | sgRNA sequence |
|---|---|
| sgRNA control, non-targeting | 5'- GACCGCGCCAAACGTGCCCTGACGG-3' |
| sgRNA human *EIF4EBP1* #1 | 5'- GGACATGGTCTCCTGTGCGCG -3' |
| sgRNA human *EIF4EBP1* #2 | 5'- GTGCGCTGCACCCGCGAACCG -3' |
| sgRNA mouse *ACACA* | 5'- GCCCAGCACATCTCGGCGCAG -3' |

**Supplementary table 3:** List of RT-qPCR primers

| Primers target gene | sequence |
|---|---|
| *ACACA* | FW: 5'- GCTGGTCCACATGAACAGG -3'<br>RV: 5'- GCCTTCTGGATATTCAGGACTTT -3' |
| *EAP1* | FW: 5'- CAGCCGCTACTCACAAATC -3'<br>RV: 5'- GCTTTCTTTATTGTTACCGCTC -3' |
| *ACT1* | FW: 5'- CCAGAAGCTTTGTTCCATCC -3'<br>RV: 5'- CGGACATAACGATGTTACCG -3' |
| *EIF4EBP1* | FW: 5'- AGCCCTTCCAGTGATGAGC -3'<br>RV: 5'- TGTCCATCTCAAACTGTGACTCTT -3' |
| *Eif4ebp1* | FW: 5'- CTAGCCCTACCAGCGATGAG -3'<br>RV: 5'- CCTGGTATGAGGCCTGAATG -3' |
| GusB | FW: 5'- GTTTTTGATCCAGACCCAGATG -3'<br>RV: 5'- GCCCATTATTCAGAGCGAGTA -3' |
| PPIA | FW: 5'- TTATTTGGGTTGCTCCCTTC -3'<br>RV: 5'- AAGTGTGCCAAATCTGCAAG -3' |
| β-actin | FW: 5'- TCCCCCAACTTGAGATGTATG -3'<br>RV: 5'- ACTGGTCTCAAGTCAGTGTACAGG -3' |
| L32 | FW: 5'- GCACACTGACTACAGCCTTGA -3' |

| | |
|---|---|
| | RV: 5'- TACCCAGGTTTGGAGGTGTG -3' |

**Supplementary table 4**: List of shRNA sequences

| shRNA label and target gene | shRNA sequence |
|---|---|
| scramble shRNA, for Tet-pLKO-puro | 5'- CCGGTCCTAAGGTTAAGTCGCCCTCGCTCG AGCGAGGGCGACTTAACCTTAGGTTTTTG -3' |
| human *EIF4EBP1* shRNA#1, for Tet-pLKO-puro | 5'- CCGGGCCAGGCCTTATGAAAGTGATCTCGA GATCACTTTCATAAGGCCTGGCTTTTTG -3' |
| human *EIF4EBP1* shRNA#2, for Tet-pLKO-puro | 5'- CCGGCGGTGAAGAGTCACAGTTTGACTCGA GTCAAACTGTGACTCTTCACCGTTTTTG -3' |
| mouse *ACACA* shRNA, for pLKO.1-neo | 5'- CCGGAGATAGAATCATCGAGTTTCTCGAGA AACTCGATGATTCTATCTTTTTTG -3' |

### KEY RESOURCES TABLE

| REAGENT or RESOURCE | SOURCE | IDENTIFIER |
|---|---|---|
| Antibodies | | |
| 4E-BP1 | Cell Signaling Technology | Cat#9644; RRID: AB_2097841 |
| 4E-BP2 | Cell Signaling Technology | Cat#2845; RRID: AB_10699019 |
| ACC1 (Acetyl-CoA Carboxylase 1) | Cell Signaling Technology | Cat#4190; RRID: AB_796746 |
| ACC2 | Cell Signaling Technology | Cat#8578; RRID: AB_10949898 |
| Anti-mouse IgG | Cell Signaling Technology | Cat#7076; RRID: AB_330924 |
| Anti-rabbit IgG | Cell Signaling Technology | Cat#7074; RRID: AB_2099233 |
| Anti-β-Actin | Sigma Aldrich | Cat#A2228; RRID: AB_476697 |
| GAPDH | Cell Signaling Technology | Cat#2118; RRID: AB_561053 |
| LC3B | Cell Signaling Technology | Cat#2775; RRID: AB_915950 |
| Phospho-Acetyl-CoA Carboxylase (S79) | Cell Signaling Technology | Cat#3661; RRID: AB_330337 |
| Phospho-AMPKalpha (T172) | Cell Signaling Technology | Cat#2535; RRID: AB_331250 |
| Phospho-S6 Ribosomal Protein (S240/244) | Cell Signaling Technology | CAT#2215; RRID: AB_331682 |

| | | |
|---|---|---|
| Vinculin | Cell Signaling Technology | Cat#4650; RRID: AB_10559207 |
| Phospho-ULK1 (S555) | Cell Signaling Technology | Cat#5869; RRID: AB_10707365 |
| eIF4E | Cell Signaling Technology | Cat#9742; RRID: AB_823488 |
| FASN (Fatty acid synthase) | Cell Signaling Technology | Cat#3180; RRID: AB_2100796 |
| ACLY | Cell Signaling Technology | Cat#13390; RRID: AB_2798203 |
| AMPKalpha | Cell Signaling Technology | Cat#2532; RRID: AB_330331 |
| IRDye® 800CW Goat anti-Mouse IgG Secondary Antibody | LI-COR Bioscience | Cat#925-32210; RRID: AB_2687825 |
| IRDye® 800CW Goat anti-Rabbit IgG Secondary Antibody | LI-COR Bioscience | Cat#925-32211; RRID: AB_2651127 |
| ULK1 | Cell Signaling Technology | Cat#8054; RRID: AB_11178668 |
| HSC-70 | Santa Cruz | Cat#sc-7298; RRID: AB_627761 |
| Mouse anti-HA-tag (F-7) | Santa Cruz | Cat#sc-7392; RRID: AB_2894930 |
| Streptavidin-HRP | ABCAM | Cat#ab59653 |
| Bacterial and Virus Strains | | |
| Competent *E.coli* Stbl3 | This study | N/A |
| Chemicals, Peptides, and Recombinant Proteins | | |
| N-acetyl-cysteine (NAC) | Sigma-Aldrich | Cat#A7250 |
| Catalase-polyethylene glycol (CAT) | Sigma-Aldrich | Cat#C4963 |
| 6-hydroxy-2,5,7,8-tetramethylchroman-2-carboxylic acid (Trolox) | Sigma-Aldrich | Cat#238813 |
| 5-(Tetradecyloxy)-2-furoic acid (TOFA) | Sigma-Aldrich | Cat#T6575 |
| Dimethyl sulfoxide (DMSO) | Sigma-Aldrich | Cat#D8418 |
| Cycloheximide (CHX) | Sigma-Aldrich | Cat#01810 |
| Crystal violet | Sigma-Aldrich | Cat#C0775 |
| Doxycycline hydrochloride (DOX) | Santa Cruz | Cat#sc-337691 |
| Critical Commercial Assays | | |
| Pierce™ BCA Protein Assay Kit | ThermoFisher Scientific | Cat#23225 |

| | | |
|---|---|---|
| NADP/NADPH-Glo™ | Promega | Cat#G9081 |
| GSH/GSSG-Glo™ | Promega | Cat#V6611 |
| Click-iT™ Protein Reaction Buffer Kit | ThermoFisher Scientific | Cat#C10276 |
| Click-iT™ EdU Cell Proliferation Kit for Imaging, Alexa Fluor™ 488 | ThermoFisher Scientific | Cat#C10337 |
| Dual-Luciferase Reporter Assay System | Promega | Cat#E1980 |
| Deposited Data | | |
| CPTAC GBM proteomic data | (Wang et al., 2021) | https://cptac-data-portal.georgetown.edu/cptac/s/S048 |
| TCGA GBM proteomic data | (Brennan et al., 2013) | |
| **From $R^2$AMC:** | | http://r2.amc.nl |
| FRENCH cohort | (Gravendeel et al., 2009) | GEO ID: gse106011 |
| TCGA bladder carcinoma | | ID: BLCA |
| TCGA Kidney renal clear cell carcinoma | | ID: KIRC |
| **From the GEPIA website:** | | http://gepia.cancer-pku.cn |
| GTEx across datasets | (Tang et al., 2017) | |
| TCGA across datasets | (Tang et al., 2017) | |
| **From the Chinese Glioma Genome Atlas website:** | | www.cgga.org.cn |
| CGGA cohort | (Zhao et al., 2021) | |
| Experimental Models: Cell Lines | | |
| Human: HEK293 (human embryonic kidney) | American Type Culture Collection (ATCC) | Cat#CRL-1573 |
| Human: HEK293-T (SV40 T-antigen containing human embryonic kidney cells) | ATCC | Cat#CRL-3216 |
| Human: U-87 MG (glioblastoma) | ATCC | Cat#HTB-14 |
| Human: HeLa (cervical adenocarcinoma) | ATCC | Cat#CRM-CCL-2 |
| Human: MCF7 (breast cancer) | ATCC | Cat#HTB-22 |
| Human: IMR-32 (neuroblastoma) | Alexander Schramm (University Hospital Essen) | N/A |
| Human: Kelly (neuroblastoma) | Alexander Schramm (University Hospital Essen) | N/A |

| | | |
|---|---|---|
| Human: Med8a (medulloblastoma) | Pablo Landgraf (University Hospital Cologne, Cologne) | N/A |
| Human: HD-MB03 (medulloblastoma) | Till Milde (DKFZ, Heidelberg) | N/A |
| Human: iPSC | TakaraBio | Cat#Y00270 |
| Human: HEK293 shRNA control (shScr) | (Dowling et al., 2010) | N/A |
| Human: HEK293 shRNAs 4EBP1, 4EBP2 (sh4EBP1/2) | (Dowling et al., 2010) | N/A |
| Mouse: MEF WT (p53-/-) and (mouse embryonic fibroblast, *Tp53* null) | (Dowling et al., 2010) | N/A |
| MEF 4EBP1/4EBP2 double knockout (DKO) (p53-/-) (*Eif4ebp1*, *Eif4ep2*, *Tp53* null) | (Dowling et al., 2010) | N/A |
| Mouse: NMuMG-NT2197 control | (Hulea et al., 2018) | N/A |
| Mouse: NMuMG-NT2197 4EBP1/4EBP2 double knockout (DKO) | (Hulea et al., 2018) | N/A |
| Mouse: GL-261 | Reuven Stein (Tel Aviv University; Israel) | N/A |
| Mouse: NIH 3T3 K-Ras$^{V12}$ | (Leprivier et al., 2013) | N/A |
| Experimental Models: Organisms/Strains | | |
| *S. cerevisiae*: BY4742 WT MATα his3Δ1 leu2Δ0 lys2Δ0 ura3Δ0 | Andreas Reichert (Heinrich Heine University, Düsseldorf) | N/A |
| *S. cerevisiae*: Eap1Δ MATα his3Δ1 leu2Δ0 lys2Δ0 ura3Δ0 eap1Δ::NatMX4 | This study | N/A |
| *S. cerevisiae*: Caf20Δ MATα his3Δ1 leu2Δ0 lys2Δ0 ura3Δ0 caf20Δ::KanMX4 | This study | N/A |
| *S. cerevisiae*: Eap1Δ/Caf20Δ MATα his3Δ1 leu2Δ0 lys2Δ0 ura3Δ0 eap1Δ::NatMX4 caf20Δ::KanMX4 | This study | N/A |
| Mouse: NOD *SCID* Gamma *Prkdc*$^{scid}$ | The Jackson Laboratory | RRID:IMSR_JAX:001303 |
| Mouse: C57BL/6J | The Jackson Laboratory | RRID:IMSR_JAX:000664 |
| Oligonucleotides | | |
| siGENOME Non-Targeting (scramble) Pool#1 | Dharmacon – Horizon Discovery | Cat#D1206-13-05 |
| siGENOME mouse *Eif4ebp1* siRNA Pool | Dharmacon – Horizon Discovery | Cat#D-05861-01 |
| siGENOME mouse *Eif4ebp2* siRNA Pool | Dharmacon – Horizon Discovery | Cat#D-044972-01 |

| | | |
|---|---|---|
| siGENOME human and mouse *EIF4E* siRNA#1 | Dharmacon – Horizon Discovery | Cat#D-003884-02 |
| siGENOME humanand mouse *EIF4E* siRNA#2 | Dharmacon – Horizon Discovery | Cat#D-003884-03 |
| siGENOME human *ACACA* siRNA#1 | Dharmacon – Horizon Discovery | Cat#D-004551-03 |
| siGENOME human *ACACA* siRNA#2 | Dharmacon – Horizon Discovery | Cat# D-004551-05 |
| siGENOME mouse *Acaca* siRNA#1 | Dharmacon – Horizon Discovery | Cat#D-063938-18 |
| siGENOME mouse *Acaca* siRNA#2 | Dharmacon – Horizon Discovery | Cat#D-063938-20 |
| ON-TARGETplus human *FASN* siRNA#1 | Dharmacon – Horizon Discovery | Cat# J-003954-12-0005 |
| ON-TARGETplus human *FASN* siRNA#2 | Dharmacon – Horizon Discovery | Cat# J-003954-14-0005 |
| mouse *Fasn* siRNA#1 | Sigma Aldrich | Cat#SASI_Mm01_00177854 |
| mouse *Fasn* siRNA#2 | Sigma Aldrich | Cat#SASI_Mm01_00177855 |
| SiRNA sequences, see Table S1 | | |
| CRISPRi sgRNA Control | This study | N/A |
| CRISPRi sgRNA targeting human *EIF4EBP1* #1 | This study | N/A |
| CRISPRi sgRNA targeting human *EIF4EBP1* #2 | This study | N/A |
| CRISPRi sgRNA targeting mouse *Acaca* | This study | N/A |
| SgRNA sequences, see Table S2 | | |
| Primers for qRT-PCR, see Table S3 | | |
| Recombinant DNA | | |
| Scramble: pLKO.1-puro scramble shRNA | Addgene | Cat#1864 |
| sh4EBP1#1: *EIF4EBP1* MISSION shRNA (human, pLKO.1-puro) | Sigma Aldrich | TRCN0000040203 |
| sh4EBP1#2: *EIF4EBP1* MISSION shRNA (human, pLKO.1-puro) | Sigma Aldrich | TRCN0000298904 |
| sh4EBP1#1: *Eif4ebp1* MISSION shRNA (mouse, pLKO.1-puro) | Sigma Aldrich | TRCN0000075610 |
| sh4EBP1#2: *Eif4ebp1* MISSION shRNA (mouse, pLKO.1-puro) | Sigma Aldrich | TRCN0000348615 |
| pLKO.1-neo | Addgene | Cat#13425 |
| shGFP: GFP shRNA (pLKO.1-neo) | Addgene | Cat#72571 |

| | | |
|---|---|---|
| shAcaca: *Acaca* shRNA (mouse, pLKO.1-neo) | This study | N/A |
| Tet-pLKO-puro | Addgene | Cat#21915 |
| Scramble inducible: Tet-pLKO-puro Non-Mammalian shRNA Control | This study | N/A |
| ish4EBP1#1: *EIF4EBP1* shRNA#1 (human, Tet-pLKO-puro) | This study | N/A |
| ish4EBP1#2: *EIF4EBP1* shRNA#2 (human, Tet-pLKO-puro) | This study | N/A |
| gRNA-dCas9-KRAB GFP | Addgene | Cat#71237 |
| sgCtrl: negative control sgRNA (gRNA-dCas9-KRAB GFP) | This study | N/A |
| sgAcaca: *Acaca* sgRNA (mouse, gRNA-dCas9-KRAB GFP) | This study | N/A |
| sg4EBP1#1: *EIF4EBP1* sgRNA#1 (human, gRNA-dCas9-KRAB GFP) | This study | N/A |
| sg4EBP1#2: *EIF4EBP1* sgRNA#2 (human, gRNA-dCas9-KRAB GFP) | This study | N/A |
| pLJM1 | Addgene | Cat#91980 |
| pLJM1-4EBP1 (T37A/T46A) [$4EBP1^{AA}$] | This study | N/A |
| pLJM1-4EBP1AA (Y54A/L59A) [$4EBP1^{AA, YL}$] | This study | N/A |
| pMSCV puro | Clontech | Cat#K1062-1 |
| pMSCV puro-4EBP1 (T37A/T46A) [$4EBP1^{AA}$] | (Goldsmith et al., 2006) | N/A |
| psPAX2 | Addgene | Cat#12260 |
| pMD2.G | Addgene | Cat#12259 |
| pGL3 control | Promega | Cat#E1741 |
| pGL3-ACACA 5'UTR (5'UTR of human *ACACA* isoform 3) | This study | N/A |
| pUb-ACC1-HA | This study | N/A |
| pUb-UTR-ACC1-HA | This study | N/A |
| pRL null *Renilla* Luciferase plasmid | Promega | Cat#E2271 |
| Software and Algorithms | | |
| R2 Genomic Analysis Visualization Platform | | http://r2.amc.nl |
| GEPIA website | (Tang et al., 2017) | http://gepia.cancer-pku.cn |
| GraphPad Prism version 7.04 and 8.0.2 | GraphPad Software | |
| cBioportal | (Cerami et al., 2012; Gao et al., 2013) | https://www.cbioportal.org/ |
| Other | | |

| | | |
|---|---|---|
| Calfectin™ Mammalian Cell Transfection Reagent | SignaGen | Cat#SL100478 |
| siLentFect™ Lipid Reagent for RNAi | BioRad | Cat#1703362 |
| CM-H2DCFDA | ThermoFisher Scientific | Cat#C6827 |
| 5'-Ethynyl-2'-deoxyuridine (EdU) | ThermoFisher Scientific | Cat#A10044 |
| Azidohomoalanine (AHA) | ThermoFisher Scientific | Cat#C10102 |
| [1-$^{14}$C]-acetate | Perkin Elmer | Cat#NEC084A001MC |

### EXPERIMENTAL MODEL AND SUBJECT DETAILS

#### Cell culture

Cells were maintained using standard tissue culture procedures in a humidified incubator at 37°C with 5% $CO_2$ and atmospheric oxygen. Stable HEK293 (human, female) control (shScr) and knock down for 4EBP1/4EBP2 (sh4EBP1/2) cell lines, WT (p53$^{-/-}$) and 4EBP1/4EBP2 double knockout (DKO) (p53$^{-/-}$) MEFs (mouse, sex unspecified) were kind gifts from Prof. Nahum Sonenberg (McGill University, Canada). NMuMG-NT2197 (mouse, female) (NT2197) control and 4EBP1/4EBP2 DKO cell lines were kind gifts from Prof. Ivan Topisirovic (McGill University, Canada). GL-261 (mouse) glioma cell line was a kind gift from Prof. Reuven Stein (Tel Aviv University, Israel). NIH 3T3 cells (mouse, male) stably expressing K-Ras$^{V12}$ have been previously described (Leprivier et al., 2013). Wild type HEK293, HEK293-T (human, female), HeLa (human, female), U-87 MG (human, male), MCF7 (human, female) cell lines were originally obtained from American Type Culture Collections (ATCC), and iPSC (human, female) were obtained from Takara Bio. Kelly (human, female) and IMR-32 (human, male) cells lines were generously donated by Prof. Alexander Schramm (University Hospital Essen). Med8a (human, male) was a kind gift from Prof. Pablo Landgraf (University Hospital Cologne, Cologne), and HD-MB03 (human, male) cell line was generously donated by Prof. Till Milde (DKFZ, Heidelberg). NT2197 were cultured in Dulbecco's modified Eagle medium (DMEM) supplemented with 10% fetal bovine serum (FBS), 1% penicillin/streptomycin (pen/strep), 10 µg/ml insulin, and 20 mM HEPES, pH 7.5. NIH 3T3 K-Ras$^{V12}$ were cultured in DMEM supplemented with 10% bovine calf serum. HD-MB03, Kelly and IMR-32 cell lines were cultured in Roswell Park Memorial Institute (RPMI) medium supplemented with 10% FBS and 1% pen/strep. iPSC line was cultured in Biolaminin 521 LN (Biolamina AB)-coated plates containing mTeSR1 medium (Stem Cell Technologies). All other cell lines were maintained in

DMEM supplemented with 10% FBS, 1% pen/strep. All cell lines were routinely confirmed to be mycoplasma-free using Venor®GeM Classic kit (Minerva Biolabs, Berlin, Germany). All human cell lines were authenticated by STR-profiling (Genomics and Transcriptomics Laboratory, Heinrich-Heine University, Germany).

**Yeast culture**

Yeast strains (all isogenic to BY4742) were grown in complex medium containing 1% (w/v) yeast extract and 2% (w/v) peptone without (YP) or with (YPD) 2% glucose. To pour solid agar plates, 2% agar was added to medium. For dot spot assays, yeast strains were grown to an OD600 of approximately 1, washed and diluted in a series of fivefold dilutions before eventually being stamped on the corresponding agar plate before incubation at 30°C or 37°C for 3-5 days. For incubation in liquid complete YPD or glucose-free YP medium, suspensions were adjusted to an OD600 of 0.1 prior to incubation at 200 rpm at 30°C. The OD600 was measured throughout the experiment with a spectrophotometer. For survival analysis, BY4742 control or *eap1Δ* yeast strains were incubated in liquid YP medium at an OD600 of 0.1 for 2 weeks at 30°C shaking at 300 rpm prior to streaking serial dilutions onto complete YPD agar plates.

**Animal models**

All mouse work was performed in accordance with the institutional animal care use committee and relevant guidelines at the Ben-Gurion University, with protocols 34-06-2016, 35-06-2016 and 59-08-2019E. C57BL/6J (C57WT) and NOD *SCID* gamma *Prkdc$^{scid}$* mice were used. Both male and female mice from 5-8 weeks of age were used for all experiments in this study. In a specific experiment all mice were from the same sex and same age. All mice were housed under specific-pathogen-free (SPF) condition at the Ben-Gurion University facility.

**Xenograft tumor models**

For sub-cutaneous injection, cancer cells ($5x10^6$-$1x10^7$) were injected into the flank or mammary fat pad of mice. Tumors size was monitored using calipers. When tumors reached the maximum of allowed size, mice were sacrificed, tumors were excised and weighed. Each tumor was cut in half and either fixed in formaldehyde 4% or snap-frozen in liquid nitrogen. When an inducible system were used, mice received 10 mg/kg/day, such that 0.05 mg/mL of doxycycline was added to the drinking water twice a week.

For orthotropic/intracranial injection, cancer cells were engrafted into the mouse brain using a stereotactic device. At the end of the experiment, mice were sacrificed and their brains were excised.

### METHOD DETAILS

#### Reagents

Cycloheximide (CHX), N-acetylcysteine (NAC), Catalase (CAT), 5-(Tetradecyloxy)-2-furoic acid (TOFA), 6-hydroxy-2,5,7,8-tetramethylchroman-2-carboxylic acid (Trolox) were from Sigma-Aldrich. Doxycycline hydrochloride (DOX) was from Santa Cruz.

#### Glucose and amino acids starvation of cell culture

Glucose or amino acids starvation was performed with subconfluent cultures (~50% confluency). For glucose starvation, full medium was replaced with DMEM or RPMI containing no glucose and no sodium pyruvate supplemented with 10% dialyzed FBS and 1 mM glucose. For amino acid starvation, full medium was replaced with Earle's Balanced Salt Solution (EBSS) supplemented with 10% dialyzed FBS and 25 mM glucose. When indicated, cells were treated with either CHX (2 µg/ml), NAC (3 mM), Catalase (400 U/ml), TOFA (5 µM) at the time of medium replacement.

#### Vectors for genetically manipulating cell lines

#### shRNA expression plasmids

To generate the shRNA expression vectors that were not commercially available, complementary oligonucleotides corresponding to shRNAs targeting mouse *Acaca* were custom cloned (Genewiz) into AgeI and EcoRI restriction sites of the pLKO.1-neo (gift from Sheila Stewart [Addgene plasmid #13425; http://n2t.net/addgene:13425; RRID:Addgene_13425]). Complementary oligonucleotides corresponding to shRNAs targeting human *EIF4EBP1* were custom cloned (Genewiz) into AgeI and EcoRI restriction sites of the Tet-pLKO-puro (gift from Dmitri Wiederschain [Addgene plasmid #21915; http://n2t.net/addgene:21915; RRID:Addgene_21915]) vectors. Cloned shRNA sequences can be found in supplementary table 4. PLKO.1-puro scramble shRNA was a gift from David Sabatini (Addgene plasmid# 1864; http://n2t.net/addgene:1864; RRID:Addgene_1864) and pLKO.1-neo shGFP was a gift from Kevin Janes (Addgene plasmid# 72571; http://n2t.net/addgene:72571; RRID:Addgene_72571). All other pLKO.1 lentiviral shRNA vectors were pLKO.1-puro based and were retrieved from the arrayed Mission TRC genome-wide shRNA collections purchased from Sigma-Aldrich Corporation.

#### CRISPRi/Cas9 plasmids

To construct CRISPRi/Cas9 targeting vectors, a non-targeting control single guide RNA (sgRNA), or sgRNAs targeting human *EIF4EBP1* or mouse *Acaca* were synthesized and custom cloned (Genewiz) into BsmBI restriction site of gRNA-dCas9-KRAB GFP (gift from

Charles Gersbach [Addgene plasmid #71237; http://n2t.net/addgene:71237; RRID:Addgene_71237]). sgRNA sequences can be found in supplementary table 2.

**cDNA expression plasmids**

The cDNA sequences of human 4EBP1 (T37A/T46A) [$4EBP1^{AA}$] and $4EBP1^{AA}$ (Y54A/L59A) ($4EBP1^{AA, YL}$) were synthesized and custom cloned (Genewiz) into the EcoRI restriction site of the pLJM1 expression vector (gift from Joshua Mendell [Addgene plasmid #91980; http://n2t.net/addgene:91980; RRID:Addgene_91980]). The cDNA sequence of human *ACACA* flanked by three HA tag sequences in 3' was synthesized and assembled (Vector Builder) in a custom made bacterial vector containing a human ubiquitin C promoter, referred as pUb-ACC1-HA. The 5'UTR of human *ACACA* isoform 3 was synthesized and custom cloned (Genewiz) into the SacI restriction site of pUb-ACC1-HA vector.

**siRNA transfections**

Cells were transfected at ~25% confluency in 6-well plates with 25 nM control ON TARGET plus non-targeting siRNA (Dharmacon) or with 25 nM of single siRNAs targeting human and mouse *EIF4E*, human or mouse *FASN*, human or mouse *ACACA*, and mouse *Eif4ebp1* and *Eif4ebp2* using siLentFect transfection reagent (Bio-Rad) according to the manufacturer's instructions. When indicated, cells were glucose starved 48 hrs post-transfection. siRNA sequences can be found in supplementary table 1.

**Virus production and viral transduction of cell lines**

HEK293-T cells were transfected with expression vectors and lentiviral packaging plasmids psPAX2 (gift from Didier Trono [Addgene plasmid #12260; http://n2t.net/addgene:12260; RRID:Addgene_12260]) and pMD2.G (gift from Didier Trono [Addgene plasmid #12259; http://n2t.net/addgene:12259; RRID:Addgene_12259]) in a ratio of 4:3:1 using CalFectin transfection reagent (Signagen) according to the manufacturer's guidelines. Medium was harvested 72 hrs post-transfection, passed through a 0.45 µm nitrocellulose filter and frozen at -80°C. Recipient cells were seeded in 6-well plates and were infected the next day when reaching ~50% confluency. For infection, 0.3 ml of virus-containing medium was added to each well in a final volume of 2 mL medium containing 8 µg/ml polybrene. Stable cell lines were either selected with 2 µg/ml puromycin or 1 mg/ml G418, or FACS sorted for cells expressing GFP.

**Immunoblot analyses of protein expression**

Cells were lysed in RIPA buffer (150 mM NaCl, 50 mM Tris-HCl, pH 8, 1% Triton X-100, 0.5% sodium deoxycholate, and 0.1% SDS) supplemented with cOmplete™, EDTA-free Protease

Inhibitor Cocktail (Sigma) and phosphatase inhibitors (PhosphoSTOP, Roche). Cell lysates were centrifuged at 14,000 x g for 15 min at 4°C and supernatants were collected. Protein concentration was measured using the Pierce™ BCA Protein Assay Kit (Thermo Fisher Scientific) according to manufacturer's protocol. Protein lysates were resolved by SDS-PAGE and transferred to nitrocellulose membranes (GE Healthcare). Membranes were blocked with 5% BSA TBS-Tween (20 mM Tris-HCl, pH 7.4, 150 mM NaCl, 0.1% Tween 20) and probed with the primary antibodies indicated in the key resources table. Secondary anti-mouse (926-32210, Li-Cor) or anti-rabbit (926-32211, Li-Cor) antibodies were used and fluorescent signal was detected with the LI-COR Odyssey CLx system.

**RNA analysis**

RNA was extracted using the RNeasy mini kit (QIAgen) according to manufacturer's instruction. cDNAs were synthesized from total RNAs using either QuantiTect Reverse Transcription Kit (QIAgen) or High-Capacity cDNA Reverse Transcription Kit (Applied Biosystems) according to manufacturer's instruction. The cDNAs were quantified by real-time PCR analysis using SYBR Green Master Mix (Bio-Rad). The primer sequences are listed in supplementary table 3. As internal controls, L32 or PPIA, GusB and β-actin were amplified.

**ROS measurements**

Cells were incubated with 5 μM chloromethyl-2′,7'-dichlorodihydrofluorescein diacetate (CM-H2DCFDA) at 37°C for 20 min. Cells were harvested and resuspended in PBS. Green fluorescence intensity was measured with a CytoFLEX flow cytometer (Beckmann Coulter). Data analysis was performed with FlowJo 10 software (FlowJo).

**Reduced and oxidized glutathione measurements**

Cells were seeded into 12-well plates and allowed to attach overnight. Cells were collected and cellular concentrations of reduced and total GSH were quantified using the GSH-Glo assay kit, according to the manufacturer's protocol (Promega). Luminescence was measured using the Spark® plate reader (Tecan).

**Liquid chromatography–mass spectrometry analysis**

To measure ATP, $NADP^+$ and NADPH, metabolite extraction was performed, in a mixture ice/dry ice, by a cold two-phase methanol–water–chloroform extraction (Elia et al., 2017; van Gorsel et al., 2019). The samples were resuspended in 700 μl of precooled methanol/water (5/3) (v/v) and 100 μL of 13C yeast internal standard. Afterwards, 500 μl of precooled chloroform was added to each sample. Samples were vortexed for 10 min at 4°C and then centrifuged (max. speed, 10 min, 4°C). The methanol–water phase containing polar metabolites was separated and dried using a vacuum concentrator at 4 °C overnight and stored at −80 °C. The measurement of ATP, $NADP^+$ and NADPH was performed by liquid chromatography using high resolution and triple quadrupole mass spectrometer. For high resolution mass spectrometry, a Dionex UltiMate 3000 LC System coupled to a Q Exactive Orbitrap mass spectrometer (Thermo Scientific) operating in negative mode was used. Metabolite separation was performed at 25°C with an Ultra High Performance Liquid Chromatography (UHPLC) from Thermo Scientific on a HILIC Fusion (P) column (150 x 2.1 mm, 5 µm). Data was collected and integrated using Xcalibur software (Thermo Scientific). Alternatively, targeted measurements of polar metabolites were performed with a 1290 Infinity II HPLC (Agilent) coupled to a 6470 triple quadrupole mass spectrometer (Agilent). Samples were injected onto a iHILIC-Fusion(P) column. Data analysis was performed with the Agilent software Masshunter. Metabolite levels were normalized to a fully 13C-labelled yeast extract and protein content.

**$NADP^+$/NADPH measurements**

NADP/NADPH-Glo™ kit (Promega) was also used to measure $NADP^+$ and NADPH. Cells were lysed in a base solution (100 mM sodium carbonate, 20 mM sodium bicarbonate, 10 mM nicotinamide, 0.05% Triton X-100) containing 1% of Dodecyltrimethylammoniumbromid (DTAB). Cell lysates were split in two equal fractions. The pH of one of the fraction was adjusted by adding 0.4 N HCl according to the manufacturer's protocol. Both fractions were then heated for 15 min at 60°C and subsequently incubated at RT for 10 min. According to the manufacturer's protocol, before adding the detection reagent, Trizma base or HCl/Trizma solution were used to adjust pH each fraction. Finally, luminescence of each fraction was analyzed with Spark® plate reader (Tecan) to measure NADP+ and NADPH levels, and the $NADP^+$/NADPH ratio was calculated.

**Protein synthesis rate**

To quantify levels of newly synthesized proteins, 50 µM of azidohomoalanine (AHA) (Thermo Fisher Scientific, Massachusetts) was added to the cell culture medium and cells were incubated for 4 hrs. Cells were then washed with ice-cold PBS, collected and lysed with EDTA-free RIPA lysis buffer (150 mM NaCl, 50 mM Tris pH 8, 1% Triton X-100, 0.5% sodium deoxycholate, 0.1% SDS). The concentration of proteins was measured by bicinchoninic acid assay using Pierce™ BCA Protein Assay Kit (Thermo Fisher Scientific), and a Click reaction was performed with Click-iT® Protein Reaction Buffer Kit (Thermo Fisher Scientific) according to manufacturer's instructions.

**Cell proliferation**

To assess cell proliferation, cells plated in 6-wells were incubated in fresh medium containing 10 µM 5-ethynyl-2'-deoxyuridine (EdU) (Invitrogen) for 60 min at 37°C. EdU staining was conducted using Click-iT™ EdU Alexa Fluor™ 488 Flow Cytometry Assay Kit (Invitrogen) according to the manufacturer's protocol. Briefly, cells were harvested, fixed with 4% paraformaldehyde in phosphate buffer saline (PBS) for 15 min, and permeabilized with 1X Click-iT™ saponin-based permeabilization reagent. Cells were incubated with a Click-iT™ reaction cocktail containing Click-iT™ reaction buffer, $CuSO_4$, Alexa Fluor® 488 Azide, and reaction buffer additive for 30 min while protected from light. Green fluorescence intensity was measured with a CytoFLEX flow cytometer (Beckmann Coulter). Data analysis was performed with FlowJo 10 software (FlowJo).

**Cell death assays**

Cell death was measured by flow cytometry using propidium iodide (PI) staining. Briefly, attached and detached cells were harvested, centrifuged and resuspended in PBS containing 1 µg/ml PI (Sigma). Cell death quantification was performed using a CytoFLEX flow cytometer (Beckmann Coulter). A minimum of 50,000 events were recorded for each replicate. Data analysis was performed with FlowJo 10 software (FlowJo).

**Soft agar colony assays**

Cells were plated in 6-well plates with 8,000 cells per well in DMEM 10% FBS or DMEM 10% bovine calf serum in a top layer of 0.25% agar added over a base layer of 0.4% agar in DMEM 10% FBS or DMEM 10% bovine calf serum. Cells were fed once a week with 1 ml of corresponding medium onto the top layer. Where indicated, NAC (5 mM), Catalase (200 U/ml), Trolox (100 µM), or TOFA (10 µM) were added to the top agar layer, as well as twice per week in the feeder medium. After 2-3 weeks at 37°C, colonies were stained with 0.01% crystal violet and 10 random fields were counted manually for each well. The percentage of colony forming cells was calculated.

**$^{14}C$ labeling and fatty acid synthesis activity**

Cells were glucose starved for 24 hrs and labeled with 10 µCi of [1-$^{14}C$]-acetate (Perkin Elmer) in the last 18 hrs. Cells were snapped frozen and lipids were extracted by methanol-water-chloroform extraction. Phase separation was achieved by centrifugation at 4°C. Radioactivity in the chloroform phase containing fatty acids was quantified by liquid scintillation counting and values were normalized to protein concentration determined in the dried protein interphase.

**5'UTR Luciferase assays**

The 5’UTR Firefly Luciferase reporter plasmids were custom cloned (Genewiz) by inserting the 5’UTR of human *ACACA* isoform 3 into the SacI and BglII restriction sites of pGL3 control vector (Promega).
For transfection, HEK293 cells were seeded in 12-well plates and transfected with 250 ng of each 5’UTR Firefly Luciferase reporter and 3 ng Renilla Luciferase expressing pRL null plasmid (Promega), completed to 500 ng DNA with pcDNA3.1 plasmid, using CalFectin transfection reagent (Signagen) according to the manufacturer’s guidelines. Cells were harvested 48 hrs post-transfection and activity of Firefly and Renilla Luciferase were sequentially determined using the Dual-Luciferase Reporter Assay System (Promega) and analyzed with the Spark® plate reader (Tecan). All samples were performed in triplicate and the final luciferase quantification was formulated as the ratio of Firefly luciferase to Renilla luciferase luminescence.

**Polysome analysis**

Cells were treated with 10 µg/ml of cycloheximide for 10 min, washed twice with PBS containing 100 µg/ml cycloheximide, then cells were scrapped and collected. Cells were pelleted by centrifugation (300 x g, 5 min, 4°C), lysed with 434 µl of lysis buffer (50 mM Tris-base pH 8, 2.5 mM $MgCl_2$, 1.5 mM KCl, 115 µg/ml cycloheximide, 2.3 mM DTT and 0.27 U/µl RNaseOUT [Thermo Fisher Scientific]), and vortexed. 25 µl of 100% Triton X-100 and 25 µl of 10% sodium deoxycholate were added to the cell lysates, which were vortexed and centrifuged (17,800 x g, 2 min, 4°C). 50 µl of the lysates were saved as the total fraction and the remaining were loaded on top of a three layers sucrose gradient (5%, 34% and 55% sucrose) that were prepared by dissolving sucrose in gradient buffer (4 mM HEPES pH 7.6, 20 mM KCl, 1 mM $MgCl_2$). The lysates loaded on top of the sucrose gradient were subjected to ultracentrifugation (229,884 x g, 2.5 hrs, 4°C). The polysome profile was read using a piston gradient collector (Biocomp) fitted with a UV detector (Tirax). Three polysomal fractions were collected and placed in Trizol (Sigma-Aldrich Company, location). RNA was extracted from frozen fractions using manufacturer's instructions.

**Bioinformatics analyses of gene and protein expression patterns in human tissue samples**

For gene expression analysis, RNA-seq data from TCGA and the GTEx projects were analyzed with Gepia (Tang et al., 2017). For survival analysis, RNA-seq and microarray data were analyzed with Kaplan-Meier Plotter (Gyorffy, 2021) or by using the Chinese Glioma Genome Atlas (CGGA; (Zhao et al., 2021)). For protein expression analysis of 4EBP1 level, CPTAC GBM proteomic data were analyzed with GraphPad Prism. For co-expression analysis of 4EBP1 and ACC1 levels, CPTAC and TCGA GBM proteomic data were analyzed by cBioportal (Cerami et al., 2012; Gao et al., 2013). For details of the cohorts, see the key ressources table.

**Quantification and Statistical Analysis**

All experiments were, if not otherwise stated, independently carried out at least three times. Statistical significance was calculated using Student's t-test in GraphPad Prism 8. The data are represented as means +/- standard deviation. A p-value of less than 0.05 was considered to be significant.
